# Spatial Regulation of Lysosomal Vesicle Acidification Along the Axon via mRAVE-Dependent v-ATPase Assembly

**DOI:** 10.64898/2025.12.22.696043

**Authors:** Surbhi Verma, Nireekshit Addanki Tirumala, Xiaolin Zhu, Raffaella De Pace, Juan S. Bonifacino

**Affiliations:** Division of Neuroscience and Cellular Structure, Eunice Kennedy Shriver National Institute of Child Health and Human Development, National Institutes of Health, Bethesda, Maryland 20892, USA

## Abstract

Lysosomal acidification is essential for neuronal homeostasis, supporting degradative clearance and metabolic signaling in all neuronal domains. Yet, how lysosomal acidification is spatially regulated within neurons remains unclear. Here, we show that assembly of the membrane-embedded V_0_ and cytosolic V_1_ domains of the vacuolar H⁺-ATPase (v-ATPase) — the proton pump that drives lysosomal acidification — governs spatial and functional lysosome diversity. In non-neuronal cells, V_1_–V_0_ association is higher in perinuclear lysosomes, correlating with increased acidity of this population. In neurons, axonal V_0_-positive vesicles move bidirectionally, whereas V_1_–V_0_ -positive vesicles move almost exclusively in the retrograde direction, consistent with the higher acidity of retrograde lysosomal vesicles. Depletion of DMXL2, a subunit of the mRAVE complex that promotes V_1_–V_0_ assembly, reduces V_1_ association, acidification, transport, and proteolytic activity of retrograde lysosomal vesicles in the axon. Together, these findings reveal a spatially regulated mechanism for the acidification of axonal lysosomal vesicles and identify mRAVE-dependent v-ATPase assembly as a key determinant of this process.

Subjects: Organelles, Trafficking, Disease

## Introduction

Lysosomes are membrane-bound organelles that serve as the primary degradative compartments within the eukaryotic endomembrane system (Ballabio & Bonifacino, 2020; Saftig & Klumperman, 2009; Xu & Ren, 2015). Besides degradation, lysosomes also function as dynamic signaling hubs for intracellular nutrient sensing and signal transduction (Ballabio & Bonifacino, 2020; Lawrence & Zoncu, 2019). The degradative function of lysosomes is closely linked to its molecular composition. The lumen contains approximately 60 different acid hydrolases, including proteases, lipases, nucleases, and glycosidases, which depend on a low pH for optimal activity. The limiting membrane is enriched with proteins that maintain lysosomal integrity and regulate its function, such as lysosome-associated membrane proteins (i.e., LAMPs), as well as a diverse array of ion channels and transporters that control acidification, solute flux, and ion homeostasis.

A key driver of lysosomal acidification is the transmembrane, vacuolar-type H^+^-ATPase (v-ATPase), an ATP-dependent proton pump responsible for actively translocating protons into the lysosomal lumen and thereby maintaining its acidic pH of 4.5–5.0 (Freeman et al., 2023; Maxson & Grinstein, 2014; Mindell, 2012). The v-ATPase is a multi-subunit complex broadly divided into two domains: the peripheral V_1_ domain, responsible for ATP hydrolysis, and the membrane-embedded V_0_ domain, responsible for proton translocation (Oot et al., 2017; Toei et al., 2010; Vasanthakumar & Rubinstein, 2020) (Fig. 1 A). V_1_ is composed of A_3,_ B_3,_ C, D, E_3,_ F, G_3_, and H subunits, while V_0_ comprises a, c_8_, c’, c″, d, e and f subunits (copy number of each subunit indicated in subscript). Different tissues and organelles express distinct isoforms of several v-ATPase subunits, contributing to the functional heterogeneity of the pump (Oot et al., 2017; Toei et al., 2010; Vasanthakumar & Rubinstein, 2020).

**Figure 1.**
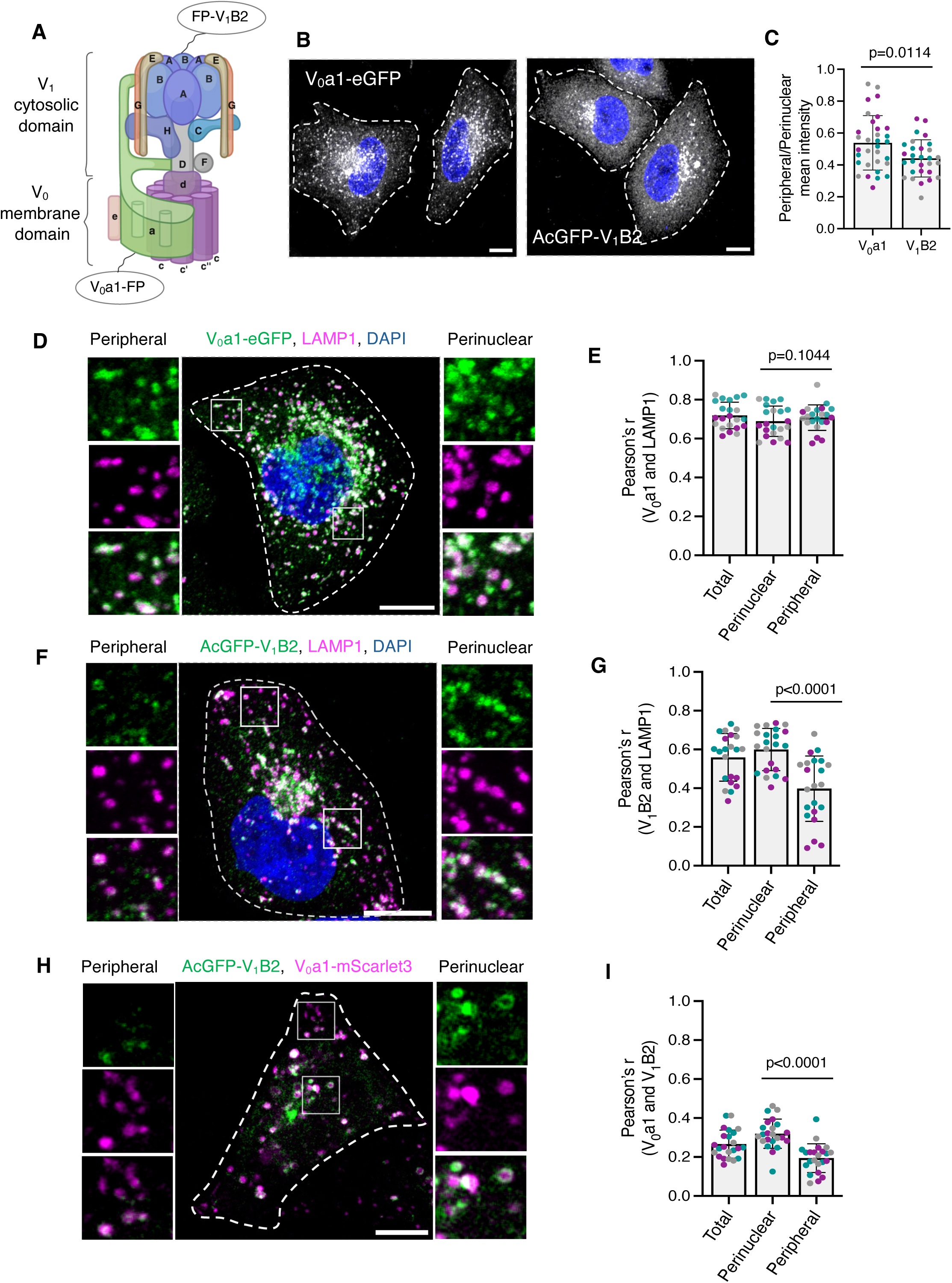
Subcellular distribution and co-localization of v-ATPase subunits V_0_a1 and V_1_B2 in HeLa cells. **(A)** Schematic representation of the v-ATPase complex highlighting the subunits analyzed in this study: V_0_a1 (membrane-embedded V_0_ domain) and V_1_B2 (cytosolic V_1_ domain). Fluorescent proteins (FP) were fused to the C-terminus of the V_0_a1 subunit and the N-terminus of the V_1_B2 subunit. **(B)** Immunofluorescence microscopy of HeLa cells stably transfected with V_0_a1–eGFP or AcGFP–V_1_B2 constructs. Single-channel images are shown in grayscale, with DAPI (nuclei) shown in blue. **(C)** Quantification of the peripheral/perinuclear mean intensity of V_0_a1–GFP and AcGFP–V_1_B2 from experiments such as that shown in panel B (n=29-32 cells from three independent experiments). **(D,F)** Immunofluorescence microscopy showing co-localization of stably expressed V_0_a1–eGFP (green) (D) or AcGFP–V_1_B2 (green) (F) with endogenous LAMP1 (magenta). Nuclei were stained with DAPI (blue). Magnified views of the boxed areas in the peripheral and perinuclear regions are shown at left and right, respectively. **(E,G)** Quantification of the co-localization between LAMP1 and V_0_a1–eGFP (E) or AcGFP–V_1_B2 (G) in total, perinuclear, and peripheral regions, expressed as Pearson’s correlation coefficient (Pearson’s r), from experiments such as those shown in panels D and F (*n*=22 cells from three independent experiments). **(H)** Live-cell images of HeLa cells showing the co-localization of transiently co-expressed V_0_a1–mScarlet3 (magenta) and AcGFP–V_1_B2 (green). Magnified views of the boxed areas in the peripheral and perinuclear regions are shown at left and right, respectively**. (I)** Quantification of Pearson’s correlation coefficients between V_0_a1–mScarlet3 and AcGFP–V_1_B2 from experiments such as that shown in panel H (*n*=23 cells from three independent experiments). All quantitative data are represented as the mean ± SD. Statistical significance was assessed using the Welch’s t-test for panel C, and Friedman test with Dunn’s multiple comparisons test for panels E, G, and I. Actual *P* values are indicated in the figure. Scale bars: 10μm.

In addition to the v-ATPase, lysosomal acidification and pH stability rely on supporting ion fluxes across the lysosomal membrane (Mindell, 2012). Several ion channels and transporters are essential for maintaining the electrochemical gradient required for sustained proton pumping (Calcraft et al., 2009; Cang et al., 2015; Kornak et al., 2001; Soyombo et al., 2006). These proteins help dissipate the transmembrane voltage generated by proton influx, facilitating efficient v-ATPase activity and stabilizing lysosomal pH.

The activity of the v-ATPase itself is tightly regulated in response to the cellular metabolic state and to signaling cues (Collins & Forgac, 2020; Kane, 2012). One key mechanism of regulation is the reversible assembly of the V_1_ and V_0_ domains. This dynamic process allows cells to tune proton-pumping activity in response to environmental conditions such as nutrient availability and cellular demand for degradation. In non-neuronal cells, signals such as glucose starvation and growth factor withdrawal have been shown to trigger v-ATPase disassembly, reducing acidification to conserve energy. The regulator of the ATPase of vacuolar and endosomal membranes (RAVE) complex (composed of DMXL1/DMXL2, WDR7, and ROGDI subunits) plays a key role in this process, promoting V_1_ recruitment to membrane-bound V_0_ and reassembly of the active V_1_–V_0_ holoenzyme (Lee et al., 2025; Nardone et al., 2025; Seol et al., 2001).

While the regulation of v-ATPase assembly by the metabolic state of the cell has been well studied, much less is known about its regulation by lysosome positioning (Pu et al., 2016). Recent work has highlighted that lysosomes are heterogeneous organelles, particularly with respect to their luminal pH (Bussi & Gutierrez, 2024). In non-neuronal cells, peripheral lysosomes are less acidic than their perinuclear counterparts, a difference that may reflect organelle maturation or local regulation of v-ATPase activity (Johnson et al., 2016; Leung et al., 2019). In neurons, this heterogeneity is especially prominent within the axon, where lysosomal vesicles characterized by the presence of LAMPs exhibit distinct acidification profiles depending on their position and direction of movement (Ferguson, 2018; Roney et al., 2022). Anterograde lysosomal vesicles are less acidic and likely represent biosynthetic, immature, or pre-degradative compartments (Gowrishankar et al., 2017; Li et al., 2024; Lie et al., 2021; Vukoja et al., 2018). In contrast, retrograde lysosomal vesicles are more acidic and may correspond to more mature lysosomes, late endosomes, amphisomes, or autolysosomes (Farfel-Becker et al., 2019; Gowrishankar et al., 2017; Lie et al., 2021; Maday et al., 2012). The mechanisms underlying this spatial control of lysosomal vesicle pH along the axon remain poorly defined.

Given the central role of v-ATPase in establishing and maintaining lysosomal pH, we hypothesized that the spatial heterogeneity of lysosomal vesicle acidification in axons could be governed by the localized assembly of the V_0_ and V_1_ domains of the v-ATPase complex. According to this hypothesis, anterograde lysosomal vesicles would be less acidic due to limited v-ATPase assembly, whereas retrograde lysosomal vesicles would become increasingly acidic as they acquire assembled v-ATPase and become degradation-competent en route back to the soma.

In this study, we test this hypothesis by analyzing the distribution and dynamics of lysosomal vesicles containing the V_0_ and V_1_ domains of the v-ATPase along the axon. We find that V_0_ exhibits greater overall co-localization with the lysosomal membrane protein LAMP1 than V_1_, supporting the existence of distinct lysosomal vesicle populations in the axon — those lacking and those containing active v-ATPase. Moreover, V_0_-positive vesicles travel bidirectionally, whereas vesicles containing V_1_ in addition to V_0_ are transported almost exclusively in the retrograde direction. Furthermore, only retrograde vesicles are labeled by the acidification marker LysoTracker and the cathepsin B activity marker Magic Red, indicating their acidic and proteolytically active nature. Finally, we show that depletion of DMXL2 (a brain-enriched subunit of the mRAVE complex) decreases the recruitment of V_1_ and the acidification and proteolytic activity of the retrograde lysosomal vesicles, leading to the accumulation of unresolved autophagosomes in the distal axon. These results demonstrate that mRAVE-complex-dependent V_1_–V_0_ assembly regulates the spatial activation of the v-ATPase and the subsequent acidification and degradative potential of lysosomal vesicles. Together, these findings reveal a novel regulatory mechanism for establishing functional heterogeneity within the axonal lysosomal system and provide new insights into the spatial control of lysosomal physiology in neurons.

## Results

### Spatial gradient of lysosomal acidification correlates with v-ATPase V_1_–V_0_ association in non-polarized cells

Previous studies showed that perinuclear lysosomes are more acidic than peripheral lysosomes in non-polarized cells (Johnson et al., 2016; Leung et al., 2019). To investigate whether this difference correlates with association of the v-ATPase V_0_ and V_1_ domains, we generated HeLa cell lines stably expressing either the V_0_a1 subunit tagged with enhanced green fluorescent protein (eGFP) at its C-terminus or the V_1_B2 subunit tagged with *Aequorea coerulescens* green fluorescent protein (AcGFP) at its N-terminus (Fig. 1 A). Whereas V_0_ is membrane-embedded, V_1_ is primarily cytosolic and can only be recruited to membranes through interaction with V_0_ (Oot et al., 2017; Toei et al., 2010; Vasanthakumar & Rubinstein, 2020). Confocal fluorescence microscopy of the fixed HeLa cells showed that V_0_a1–eGFP localized to vesicular structures that were distributed throughout the cytoplasm but were more abundant in the perinuclear region (Fig. 1, B and C). AcGFP–V_1_B2 displayed a similar vesicular pattern, although with fewer vesicles in the peripheral area and higher cytosolic fluorescence (Fig. 1, B and C). Furthermore, V_0_a1–eGFP exhibited a high degree of co-localization with the endogenous lysosomal membrane protein LAMP1 in both the perinuclear and peripheral regions (Fig. 1, D and E). In contrast, AcGFP–V_1_B2 showed less co-localization with LAMP1 in the peripheral region compared to the perinuclear region (Fig. 1, F and G). Similar patterns of V_0_a1–eGFP and AcGFP–V_1_B2 distribution relative to LAMP1 were observed in another non-polarized cell type, U2OS (Fig. S1 A-D). Additionally, live-imaging of HeLa cells stably expressing V_0_a1–eGFP and labeled with the acidic organelle marker LysoTracker (Chazotte, 2011), showed higher co-localization of V_0_a1 with LysoTracker in the perinuclear region compared to the peripheral region (Fig. S1, E and F). Finally, live-imaging of HeLa cells transiently co-transfected with V_0_a1–mScarlet3 and AcGFP–V_1_B2 constructs revealed higher co-localization of these subunits in the perinuclear region compared to the peripheral region (Fig. 1, H and I). These findings indicate that the previously demonstrated perinuclear–peripheral gradient of lysosomal acidification (Johnson et al., 2016; Leung et al., 2019) correlates with differential V_1_–V_0_ association.

### Distinct populations of V_0_- and V_1_–V_0_-containing vesicles in the axon of rat hippocampal neurons

Spatial variations in lysosomal acidification have also been observed along the axons of rat hippocampal neurons (Ferguson, 2018; Roney et al., 2022). Specifically, lysosomal vesicles moving in the anterograde direction are less acidic, likely representing biosynthetic vesicles or lysosome precursors (Gowrishankar et al., 2017; Lie et al., 2021). In contrast, retrograde lysosomal vesicles are more acidic and may correspond to mature lysosomes, late endosomes, autolysosomes, or amphisomes (Farfel-Becker et al., 2019; Gowrishankar et al., 2017; Lie et al., 2021). To examine whether these pH differences correlate with differential association of V_0_ and V_1_ in the axon, we transiently transfected day-in-vitro-5 (DIV5) rat hippocampal neurons in primary culture with V_0_a1–eGFP or AcGFP–V_1_B2 constructs (Fig. 2 A). At DIV7, neurons were fixed, permeabilized, immunostained for LAMP1 and the axon initial segment (AIS) marker ankyrin G (AnkG) (to identify the axon), and examined by confocal fluorescence microscopy. We observed that V_0_a1–eGFP and LAMP1 extensively co-localized on vesicles that were evenly distributed along the axon (Fig. 2, B and C). In contrast, AcGFP–V_1_B2 was less membrane-associated and exhibited a lower degree of co-localization with LAMP1 (Fig. 2, D and E). Co-expression of V_0_a1–eGFP and mCherry–V_1_B2 revealed that V_0_a1–positive vesicles were more numerous than V_1_B2-positive vesicles in the axon, with only a fraction of V_0_a1–positive vesicles displaying associated V_1_B2 (Fig. 2, F and G). In contrast, virtually all V_1_B2-positive vesicles co-localized with V_0_a1–eGFP (Fig. 2, F and G). Therefore, at least two populations of lysosomal vesicles exist in the axon: those containing only the V_0_ domain and those containing both the V_0_ and V_1_ domains.

**Figure 2.**
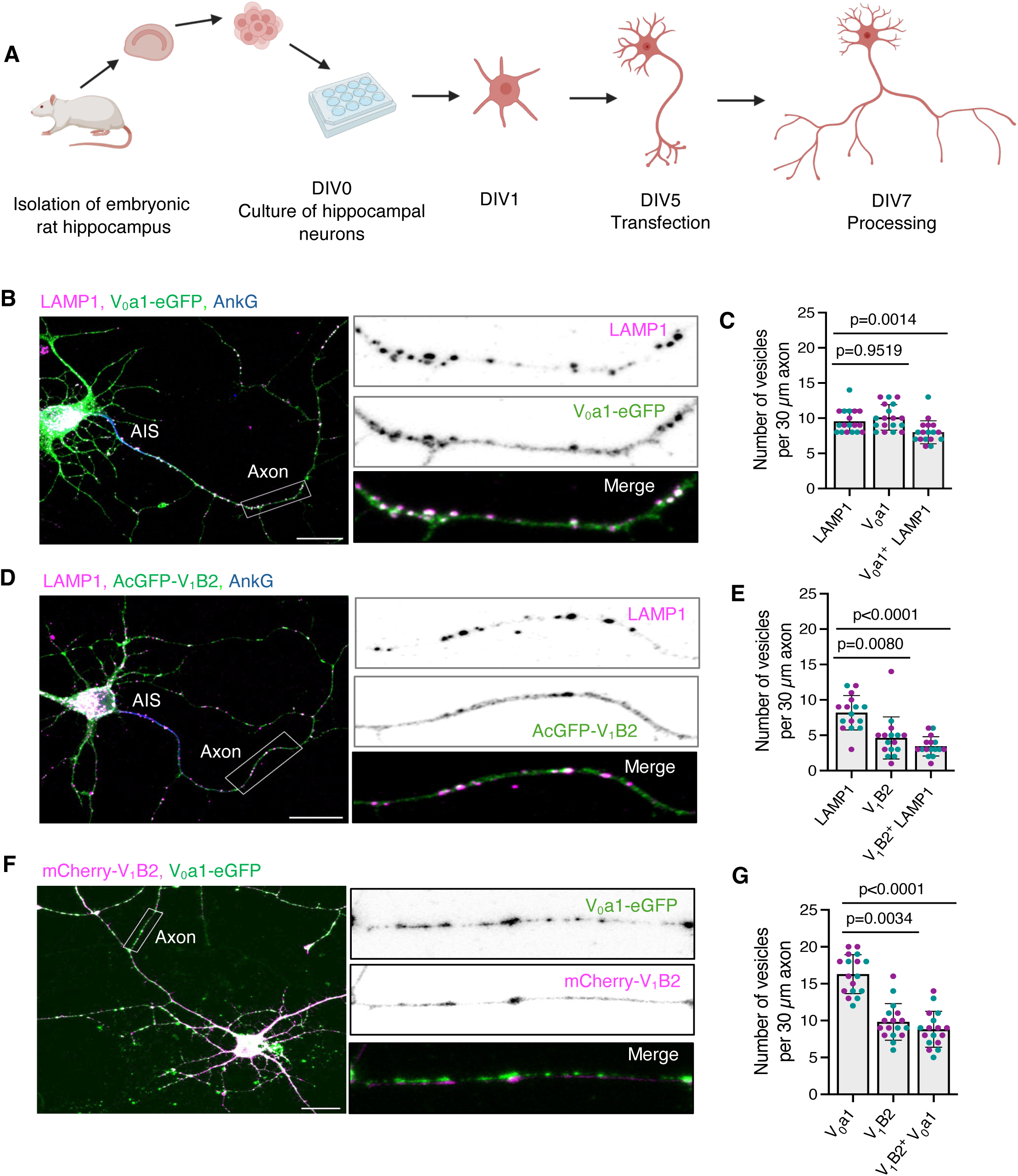
Localization of V_0_a1 and V_1_B2 subunits in axons from rat hippocampal neurons. **(A)** Schematic representation of the isolation, transfection, and processing of embryonic rat hippocampal neurons. **(B)** Immunofluorescence microscopy of DIV7 rat hippocampal neurons transiently transfected with a plasmid encoding V_0_a1–eGFP and immunostained for the endogenous lysosomal membrane protein LAMP1 (magenta) and axon initial segment (AIS) protein ankyrin G (AnkG) (blue). Magnified views of the boxed 30-μm axonal segments are shown at right. **(C)** Quantification of V_0_a1–eGFP, LAMP1, and V_0_a1–eGFP-positive LAMP1 vesicles per 30 μm axon length from experiments such as that shown in panel B (*n*=18 neurons from ≥4 cultures prepared from two rats). **(D)** Immunofluorescence microscopy of DIV7 rat hippocampal neurons transfected with a plasmid encoding AcGFP–V_1_B2 and immunostained for endogenous LAMP1 (magenta) and ankyrin G (blue). Magnified views of the boxed 30-μm axonal segments are shown at right. **(E)** Quantification of AcGFP–V_1_B2, LAMP1, and AcGFP–V_1_B2-positive LAMP1 vesicles per 30 μm axon length from experiments such as that shown in panel D (*n*=16 neurons from ≥4 cultures prepared from two rats). **(F)** Immunofluorescence microscopy of DIV7 rat hippocampal neurons co-transfected with plasmids encoding V_0_a1–eGFP and mCherry–V_1_B2. Magnified views of the boxed 30-μm axonal segments are shown at right. **(G)** Quantification of V_0_a1–eGFP, mCherry–V_1_B2, and mCherry–V_1_B2-positive V_0_a1–eGFP vesicles per 30 μm axon length from experiments such as that shown in panel F (*n*=17 neurons from ≥4 cultures prepared from two rats). All the axonal segments analyzed correspond to the mid-axon and are located approximately 40-200 μm from the soma. All quantitative data are represented as the mean ± SD. Statistical significance was assessed using the Friedman test with Dunn’s multiple comparisons test. Actual *P* values are indicated in the figure. Scale bars: 20 μm.

### V_0_-positive vesicles exhibit bidirectional movement, whereas V_1_-positive vesicles only move retrogradely in the axon

While the experiments described above demonstrated the presence of lysosomal vesicle populations containing different complements of V_0_ and V_1_–V_0_ v-ATPase forms within the axon, they did not identify the specific vesicle populations in which these forms were localized. To determine whether these v-ATPase forms were associated with lysosomal vesicles moving in different directions, we performed live-imaging of axons from rat hippocampal neurons transiently expressing either V_0_a1–mScarlet3 or mCherry–V_1_B2 at DIV7. Axons were imaged for 180 s, and vesicle dynamics were analyzed using kymographs generated from 30-μm of mid-axonal segments located approximately 40-200 μm from the soma. This analysis revealed distinct transport behaviors of vesicles containing the v-ATPase subunits. While V_0_a1–positive vesicles exhibited 31% anterograde and 47% retrograde movement (Fig. 3, A and B), V_1_B2-positive vesicles showed only 11% anterograde and 59% retrograde movement (Fig. 3, C and D) The remaining puncta were stationary. When neurons were co-transfected with V_0_a1–mScarlet3 and AcGFP–V_1_B2 constructs, imaging of mid-axonal segments revealed that among the 23% anterograde V_0_-positive vesicles, only 4% had associated V_1_, whereas among the 50% retrograde V_0_-positive vesicles, 39% had associated V_1_ (Fig. 3, E and F, see also Video S1). Imaging of V_1_ vesicles tagged with the brighter fluorescent protein mNeonGreen produced similar results, ruling out the existence of a fainter, undetected population of anterograde V_1_ vesicles and thus confirming the net retrograde movement of V_1_ (Fig. S2, A and B). In the distal axon (approximately 20-50 μm from the axon tip), a higher percentage of V_0_- and V_1_-positive vesicles were immobile, and no significant differences were observed in the motility profile of both subunits (Fig. S2, C and D). Together, these findings revealed a striking difference in the transport dynamics of V_0_ and V_1_ in the axon. While V_0_ exhibits bidirectional transport, V_1_ moves predominantly in the retrograde direction, suggesting that V_1_–V_0_ assembly occurs mainly in the distal axon, before retrograde transport toward the soma. This spatially regulated V_1_–V_0_ assembly provides a likely mechanistic basis for the differential acidification of anterograde and retrograde lysosomal vesicles in the axon.

**Figure 3.**
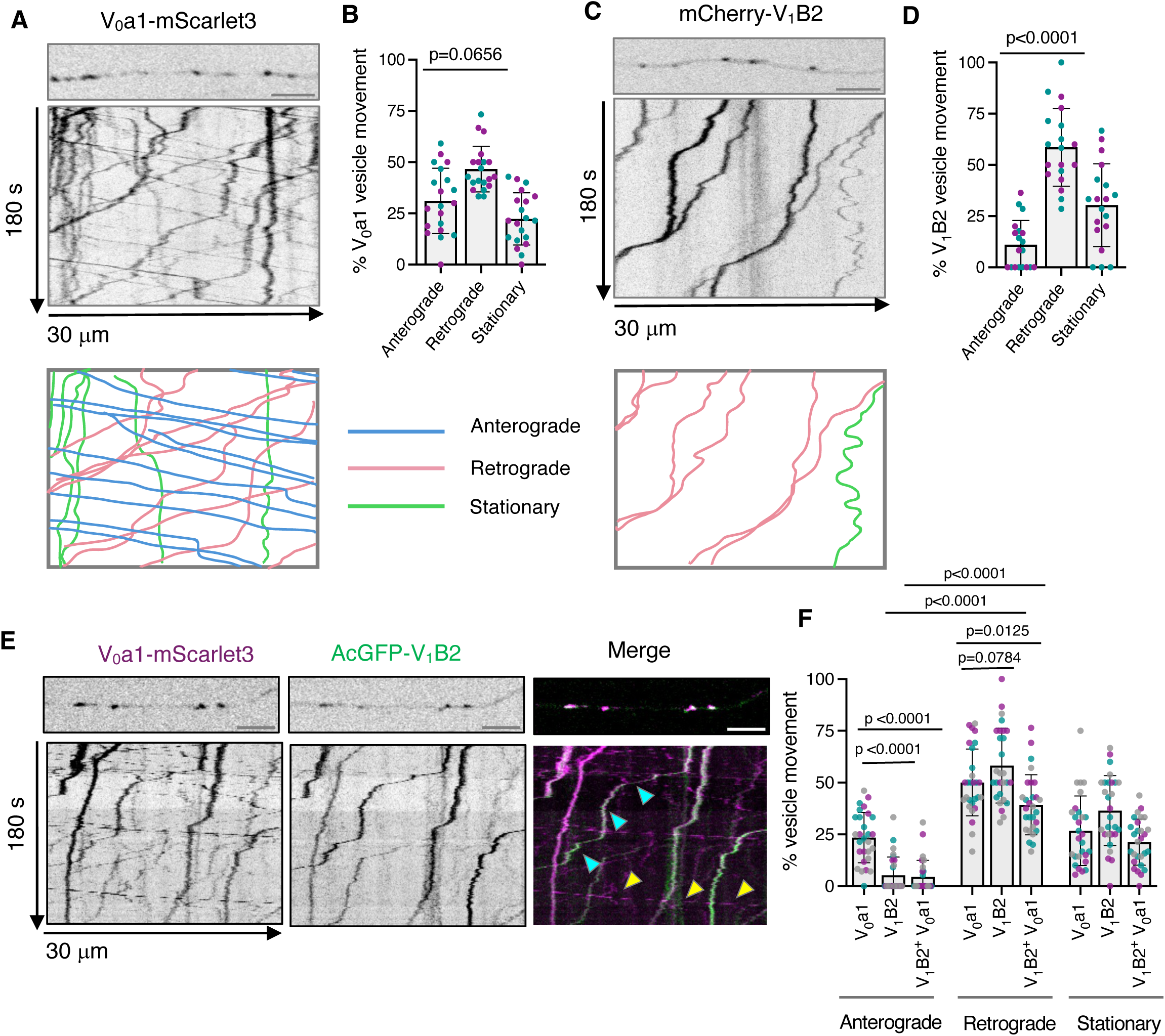
Axonal transport dynamics of vesicles containing V_0_a1 and V_1_B2 subunits in axons from rat hippocampal neurons. **(A,C)** Single frames (top) and kymographs (bottom) of 30-μm axonal segments from DIV7 rat hippocampal neurons expressing V_0_a1–mScarlet3 (A) or mCherry–V_1_B2 (C) imaged live for 180 s. The bottom schemes show representative trajectories of vesicle movement, with anterograde motion in blue (positive slopes), retrograde motion in magenta (negative slopes), and stationary vesicles in green (vertical lines). **(B,D)** Quantification of the proportion of anterograde, retrograde, and stationary vesicles in axons from neurons expressing V_0_a1–mScarlet3 (B) or mCherry–V_1_B2 (D) from experiments such as those shown in panels A and C (*n*=19-20 neurons from ≥4 cultures prepared from two rats). **(E)** Single frames (top) and kymographs (bottom) of 30-μm axonal segments from DIV7 rat hippocampal neurons co-expressing V_0_a1–mScarlet3 and AcGFP–V_1_B2 imaged live for 180 s. Cyan arrows indicate V_1_-V_0_-positive vesicles and yellow arrows indicate only V_0_-positive vesicles. **(F)** Quantification of the proportion of anterograde, retrograde, and stationary vesicles in axons from neurons co-expressing V_0_a1–mScarlet3 and AcGFP–V_1_B2 from experiments such as that shown in panel E (*n*=28 neurons from ≥4 cultures prepared from three rats). All the axonal segments analyzed correspond to the mid-axon and are located approximately 40-200 μm from the soma. All quantitative data represent the mean ± SD. Statistical significance was calculated using the Friedman test with Dunn’s multiple comparisons test for panels B and D, and using two-way ANOVA with Tukey’s multiple comparisons test for panel F. Actual *P* values are indicated in the figure. Scale bars: 5 μm.

### All V_0_ and V_1_ vesicles contain cathepsin D, but only retrograde V_0_ and V_1_ vesicles are acidic

To further investigate the nature of V_0_- and V₁-positive vesicles, we analyzed their co-transport with well-established lysosomal markers in mid-axonal segments located approximately 40-200μm from the soma of rat hippocampal neurons. One such marker, SiR-Lysosome, is a silicon–rhodamine fluorophore conjugated to pepstatin A that selectively labels the lysosomal hydrolase cathepsin D (Albrecht et al., 2020). Live imaging and kymograph analysis revealed SiR-lysosome staining in most anterograde and retrograde V_0_a1–mScarlet3-positive vesicles (Fig. 4, A and B), as well as in the retrograde mCherry–V_1_B2-positive vesicles (Fig. 4, C and D), consistent with their lysosomal identity. SiR-lysosome was also present in stationary V₀- and V₁-positive structures along the axon (Fig. 4, A-D). Additionally, live imaging of neurons co-expressing V_0_a1-mScarlet3 and mNeonGreen-V_1_B2 neurons revealed co-transport of V_0_a1 and V_1_B2 with SiR-lysosome positive vesicles (Video S2).

**Figure 4.**
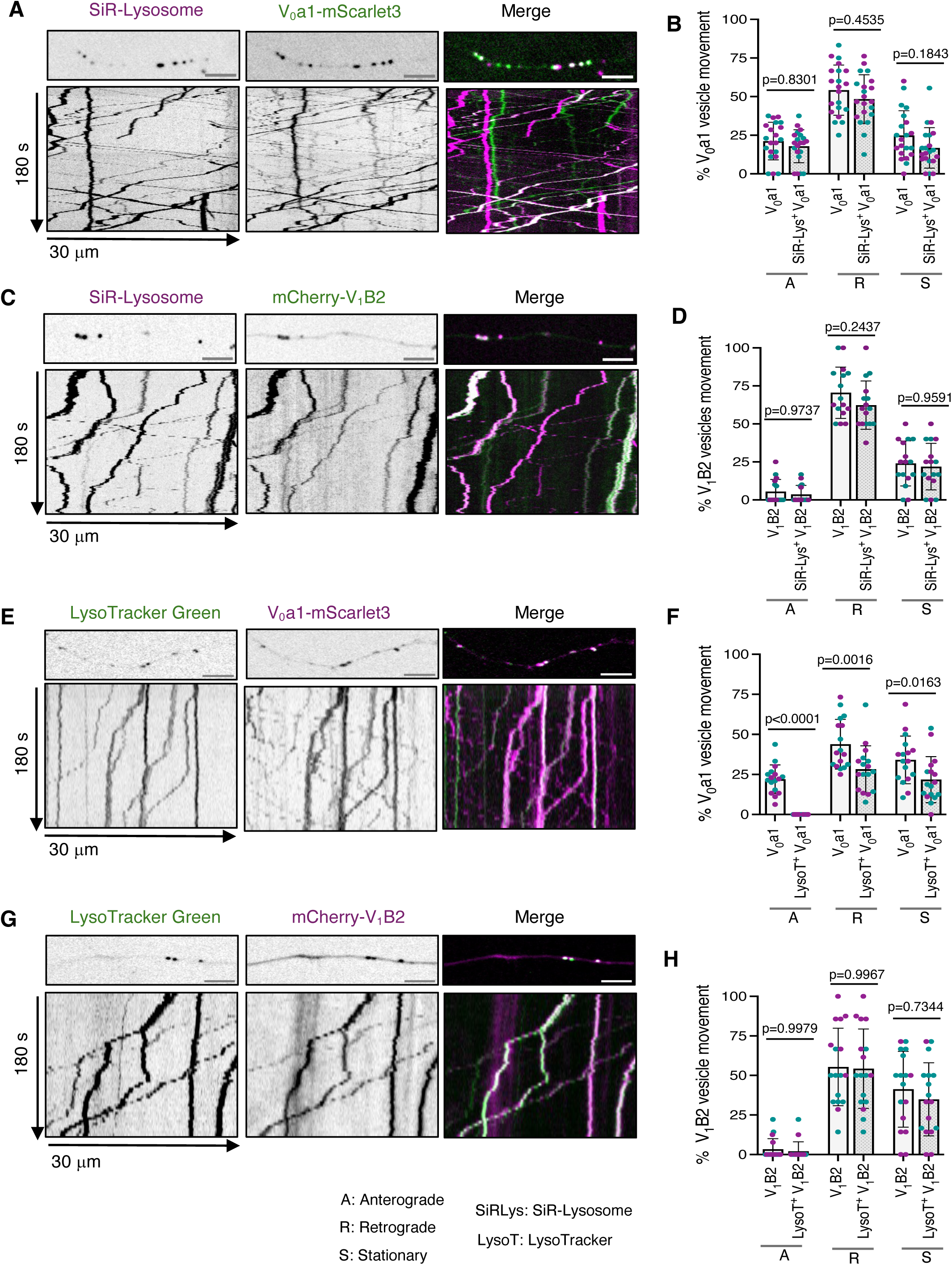
Cathepsin D content and acidification of axonal V_0_ and V_1_ vesicles in axons from rat hippocampal neurons. **(A)** Single frames (top) and kymographs (bottom) of 30-μm axonal segments from DIV7 rat hippocampal neurons transiently expressing V_0_a1–mScarlet3 and stained with the cathepsin D probe SiR-Lysosome. **(B)** Quantification of the proportion of anterograde, retrograde, and stationary V_0_a1–mScarlet3 vesicles and SiR-Lysosome-positive V_0_a1–mScarlet3 vesicles from experiments such as that shown in panel A (*n*=21 neurons from ≥4 cultures prepared from two rats). **(C)** Single frames (top) and kymographs (bottom) of 30-μm axonal segments from DIV7 rat hippocampal neurons expressing mCherry–V_1_B2 and stained with SiR-Lysosome. **(D)** Quantification of the proportion of anterograde, retrograde, and stationary mCherry–V_1_B2 vesicles and SiR-Lysosome-positive mCherry–V_1_B2 vesicles from experiments such as that shown in panel C (*n*=16 neurons from ≥4 cultures prepared from two rats). **(E)** Single frames (top) and kymographs (bottom) of 30-μm axonal segments from DIV7 rat hippocampal neurons expressing V_0_a1–mScarlet3 and stained with the acidic pH probe LysoTracker Green. **(F)** Quantification of the proportion of anterograde, retrograde, and stationary V_0_a1–mScarlet3 and Lysotracker-positive V_0_a1–mScarlet3 vesicles from experiments such as that shown in panel E (*n*=17 neurons from ≥4 cultures prepared from two rats). **(G)** Single frames (top) and kymographs (bottom) of 30-μm axonal segments from DIV7 rat hippocampal neurons expressing mCherry-V_1_B2 and stained with LysoTracker Green. **(H)** Quantification of the proportion of anterograde, retrograde, and stationary V_1_B2–mCherry vesicles and LysoTracker-positive mCherry-V_1_B2 vesicles from experiments such as that shown in panel G (*n*=17 neurons from ≥4 cultures prepared from two rats). All the axonal segments analyzed correspond to the mid-axon and are located approximately 40-200 μm from the soma. All quantitative data represent mean ± SD. Statistical significance was calculated using two-way ANOVA with Sidak’s multiple comparisons test. Actual *P* values are indicated in the figure. Scale bars: 5 μm.

To assess vesicle acidification, neurons expressing V_0_a1–mScarlet3 or mCherry–V_1_B2 were incubated with LysoTracker Green. Live imaging and kymograph analysis of the axons revealed that most LysoTracker-positive vesicles moved retrogradely, consistent with previous reports that more mature lysosomal vesicles exhibit net retrograde transport (Fig. 4, E-H). Co-localization analyses showed LysoTracker signal in most retrograde but not anterograde V_0_a1–mScarlet3-positive vesicles (Fig. 4, E and F). Likewise, retrograde mCherry–V_1_B2-positive vesicles stained for LysoTracker (Fig. 4, G and H).

Together, these experiments demonstrated that all V_0_- and V_1_-positive vesicles contained cathepsin D, regardless of their direction of transport. However, only retrograde V_0_- and V_1_-positive vesicles were labeled with LysoTracker, supporting the idea that anterograde V_0_-positive vesicles correspond to biosynthetic intermediates or lysosomal precursors, whereas only V_1_–V_0_ - positive vesicles acquire the acidity and degradative potential characteristic of mature lysosomes. These findings suggest that V_1_–V_0_ assembly in the distal axon is a key determinant of retrograde lysosomal vesicle acidification.

### Loss of DMXL2 decreases the population of retrograde V_1_–V_0_ -containing vesicles in the axon

To identify regulatory factors controlling V_1_–V_0_ assembly in the axon, we examined the effect of disrupting the metazoan RAVE (mRAVE) complex, an evolutionarily conserved regulator of v-ATPase assembly composed of DMXL1/DMXL2, WDR7, and ROGDI subunits (Jaskolka et al., 2021; Nardone et al., 2025). We depleted DMXL2 in rat hippocampal neurons by lentiviral transduction with *Dmxl2* shRNA at DIV4 and analyzed the neurons at DIV7 (Fig. 5 A). Immunoblot analysis confirmed ∼70% knockdown (KD) efficiency of DMXL2 in whole neurons using *Dmxl2* shRNA compared with a scrambled shRNA control (Fig. 5, B and C). This was accompanied by a ∼50% reduction in membrane-associated V_1_B2, but not V_0_a1, in *Dmxl2* shRNA-treated neurons (Fig. 5, D and E). In contrast, total levels of V_0_a1 or V_1_B2 in whole neuronal lysate were unchanged between control and DMXL2-KD neurons (Fig. 5, F-H).

**Figure 5.**
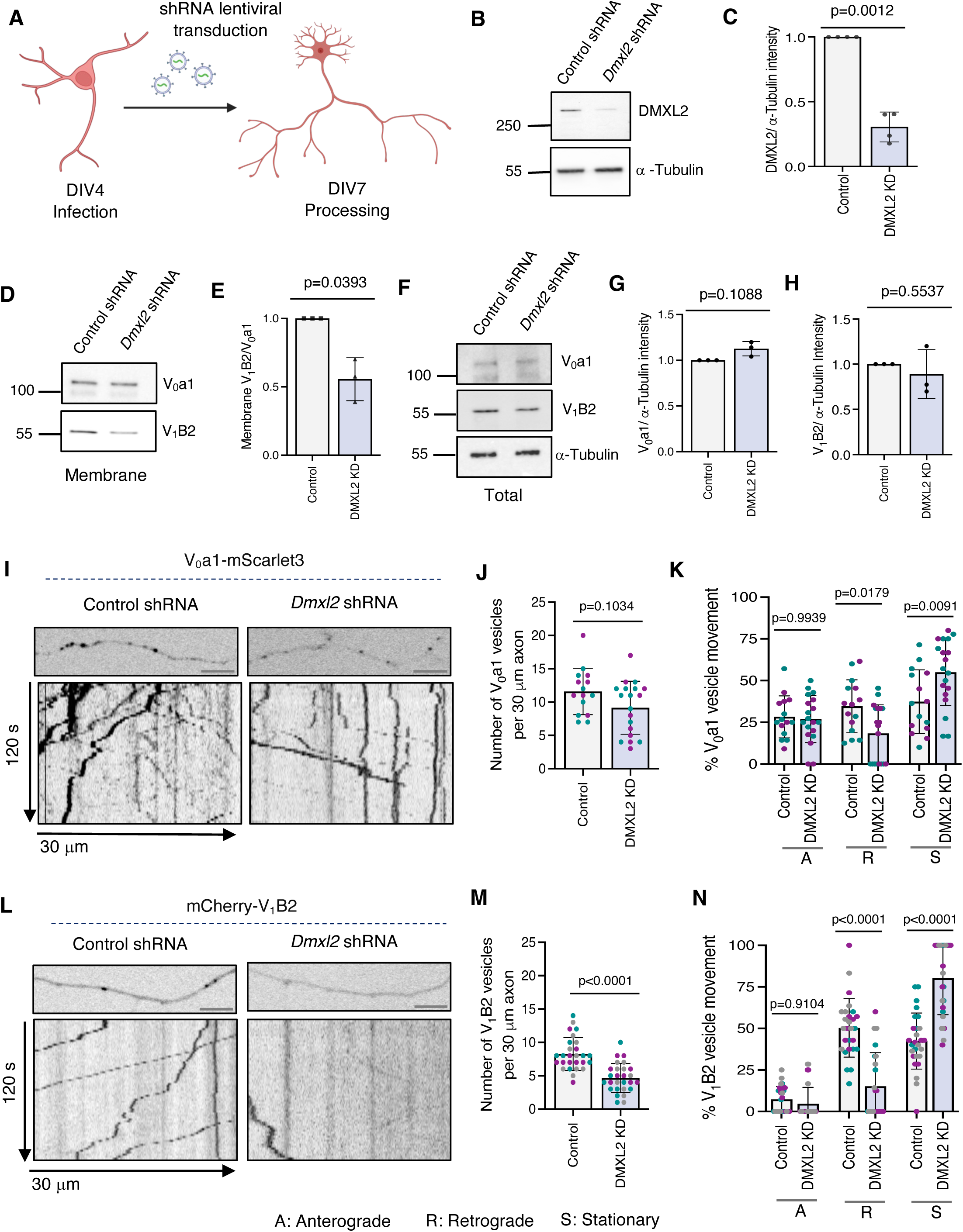
DMXL2 depletion decreases V_1_ membrane association and axonal V_0_ and V_1_ vesicle motility in rat hippocampal neurons. **(A)** Schematic representation of lentiviral shRNA transduction of rat hippocampal neurons. **(B)** SDS-PAGE and immunoblot analysis showing KD of DMXL2 using *Dmxl2* shRNA in DIV7 rat hippocampal neurons where α-Tubulin was used as a loading control. The positions of molecular mass markers (in kDa) are indicated on the left. **(C)** Quantification of immunoblots such as that shown in panel B demonstrating DMXL2-KD efficiency (*n* = 4 independent experiments). **(D)** SDS-PAGE and immunoblot analysis showing levels of V_0_a1 and V_1_B2 associated with membranes in control and DMXL2-KD rat hippocampal neurons. The positions of molecular mass markers (in kDa) are indicated on the left. **(E)** Quantification of the V_1_/V_0_ ratio in membrane fractions from control and DMXL2-KD neurons from experiments such as that shown in panel D (n=3 independent experiments). **(F)** SDS-PAGE and immunoblot analysis showing levels of V_0_a1 and V_1_B2 in control and DMXL2-KD rat hippocampal neurons, with α-Tubulin used as a loading control. The positions of molecular mass markers (in kDa) are indicated on the left. **(G-H)** Quantification of V_0_a1 (G) and V_1_B2 (H) protein levels normalized to α-Tubulin in whole-cell neuronal lysates from control and DMXL2-KD neurons, from experiments such as that shown in panel F (n=3 independent experiments). **(I)** Single frames (top) and kymographs (bottom) of 30-μm axonal segments from control and DMXL2-KD rat hippocampal neurons expressing V_0_a1–mScarlet3 imaged live for 120 s. **(J)** Quantification of the total number of V_0_a1–mScarlet3 vesicles in control and DMXL2-KD neurons from experiments such as that shown in panel I (*n*=15-19 neurons from ≥4 cultures prepared from two rats). **(K)** Quantification of the proportion of anterograde, retrograde, and stationary V_0_a1–mScarlet3 vesicles in control and DMXL2-KD neurons from experiments such as that shown in panel I (*n*=15-19 neurons from ≥4 cultures prepared from two rats). **(L)** Single frames (top) and kymographs (bottom) of 30-μm axonal segments from control and DMXL2-KD neurons expressing mCherry–V_1_B2 imaged live for 120 s. **(M)** Quantification of the total number of mCherry–V_1_B2 vesicles in control and DMXL2-KD neurons from experiments such as that shown in panel L (*n*=27 neurons from ≥4 cultures prepared from three rats). **(N)** Quantification of the proportion of anterograde, retrograde, and stationary mCherry–V_1_B2 vesicles in control and DMXL2-KD neurons from experiments such as those shown in panel L (*n*=27 neurons from ≥4 cultures prepared from three rats). Imaging was performed in DIV7 rat hippocampal neurons. All the axonal segments analyzed correspond to the mid-axon and are located approximately 40-200 μm from the soma. All quantitative data are represented as the mean ± SD. Statistical significance was calculated using Welch’s t-test for panels C, E, G and H, the Mann-Whitney test for panels J and M, and two-way ANOVA with Sidak’s multiple comparisons test for panels K and N. Actual *P* values are indicated in the figure. Scale bars: 5 μm.

Live imaging of neurons expressing V_0_a1–mScarlet3 revealed no significant change in the total number of V_0_ vesicles (Fig. 5, I and J) but showed a specific reduction in retrograde V_0_ vesicles accompanied by an increase in stationary V_0_ vesicles within the axon of DMXL2-KD neurons (Fig. 5, I and K). These findings suggest that V_1_–V_0_ assembly — and the resulting vesicle acidification — promotes the retrograde, but not the anterograde, transport of V_0_-positive lysosomal vesicles. In addition, we observed that DMXL2-KD resulted in an ∼50% reduction in the total number of mCherry–V_1_B2 vesicles in the axon (Fig. 5, L and M), likely reflecting impaired recruitment of the V_1_ domain to membrane-bound V_0_. Furthermore, DMXL2-KD caused a pronounced decrease in retrograde mCherry–V_1_B2 vesicles within the axon (Fig. 5, L and N).

From these experiments, we concluded that DMXL2: and by extension, the mRAVE complex: is a key regulator of V_1_–V_0_ assembly and of the retrograde transport of V_1_–V_0_-containing lysosomal vesicles within the axon.

### DMXL2 depletion alters the properties of lysosomal vesicles in the axon

Next, we investigated whether DMXL2 depletion alters the properties of lysosomal vesicles in the axon. To this end, we assessed lysosomal acidification and motility in rat hippocampal neurons using LysoTracker staining. We observed that DMXL2 KD decreased the total number of LysoTracker-positive vesicles in the axon (Fig. 6, A and B). Furthermore, DMXL2 KD almost completely abolished the population of retrograde LysoTracker-positive vesicles (Fig. 6, A and C). To further assess lysosomal function, we employed the Magic Red Cathepsin B probe, which fluoresces upon cleavage by active cathepsin B: a process that requires lysosomal acidification. In control neurons, Magic-Red-positive vesicles were readily detected in axons and exhibited predominantly retrograde transport (Fig. 6, D–F). In contrast, neurons transduced with *Dmxl2* shRNA displayed a pronounced reduction in both the number and retrograde motility of Magic Red–positive vesicles, indicating impaired degradative activity of axonal lysosomes (Fig. 6, D–F). Reduced Magic Red fluorescence was also observed in the soma and dendrites of DMXL2-KD neurons (Fig. 6, I and J), indicating that the effects of v-ATPase disassembly were not limited to the axon. Moreover, LC3 immunostaining revealed increased accumulation of the autophagy protein LC3 within distal axons of DMXL2-KD neurons, indicative of autophagosome accumulation (Fig. 6, G and H). This was accompanied with increased LC3 intensity in the soma of DMXL2-KD neurons (Fig. 6, K and L), also suggesting that the effects of DMXL2 KD extend beyond the axon to affect lysosomal acidification and function throughout the neuron. From these observations, we concluded that DMXL2-dependent V_1_–V_0_ assembly is critical for the acidification and proteolytic function of lysosomal vesicles throughout the neuron.

**Figure 6.**
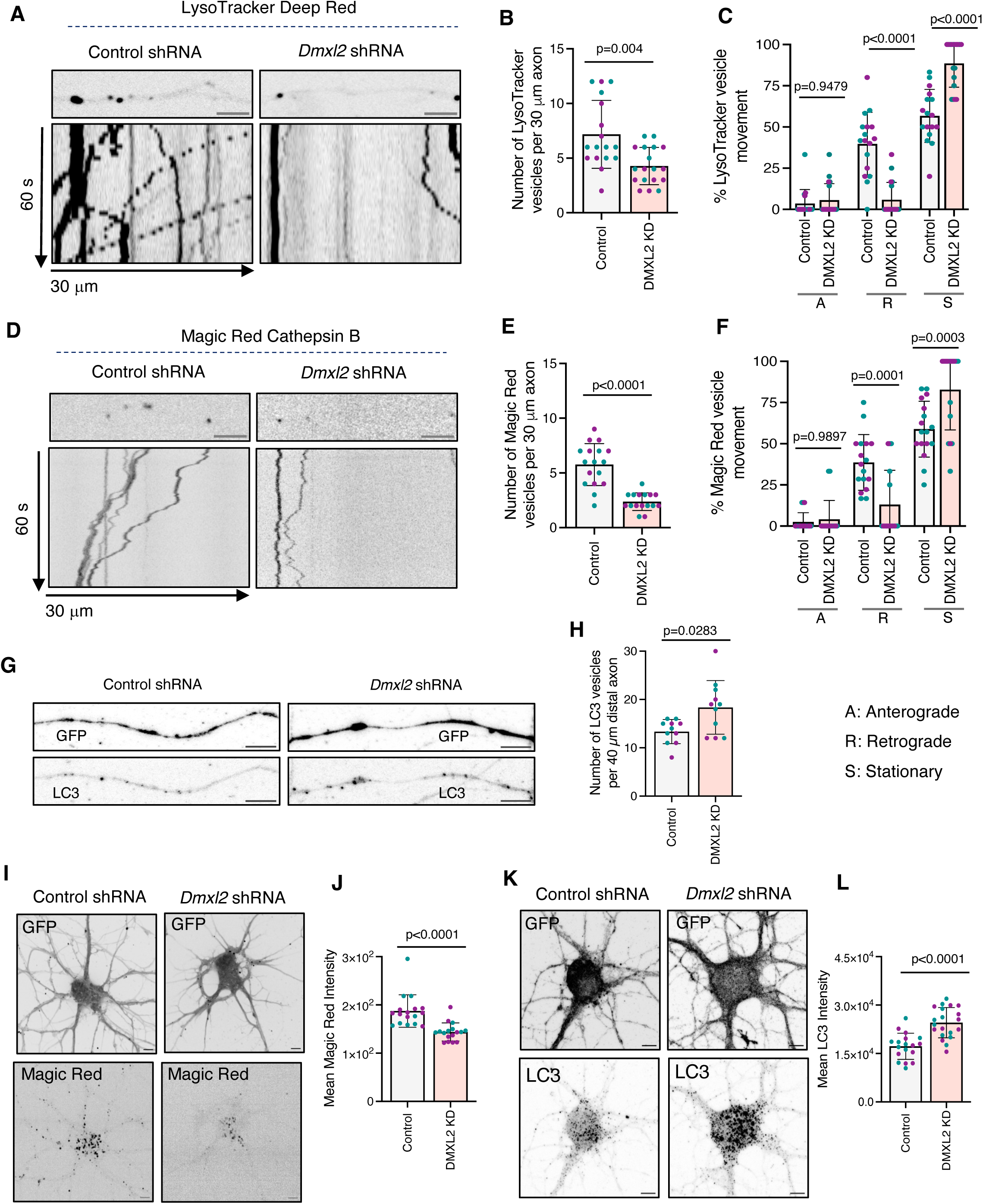
DMXL2 depletion alters lysosomal vesicle dynamics and v-ATPase subunit distribution in rat hippocampal neurons. **(A)** Single frame (top) and kymographs (bottom) of 30-μm axonal segments from control and DMXL2-KD DIV7 hippocampal neurons stained with LysoTracker Deep Red and imaged live for 60 s. **(B)** Quantification of the total number of LysoTracker-positive vesicles in control and DMXL2-KD axons from experiments such as that shown in panel A (*n*=17-18 neurons from ≥4 cultures prepared from two rats). **(C)** Quantification of the proportion of anterograde, retrograde, and stationary LysoTracker-positive vesicles in control and DMXL2-KD axons from experiments such as that shown in panel A (*n*=17-18 neurons from ≥4 cultures prepared from two rats) **(D)** Single frame (top) and kymographs (bottom) of 30-μm axonal segments from control and DMXL2-KD DIV7 hippocampal neurons stained with Magic Red cathepsin B substrate and imaged live for 60 s. **(E)** Quantification of the total number of Magic Red-positive vesicles in control and DMXL2-KD axons from experiments such as that shown in panel D (*n*=16-17 neurons from ≥4 cultures prepared from two rats). **(F)** Quantification of the proportion of anterograde, retrograde, and stationary Magic Red-positive vesicles in control and DMXL2-KD axons from experiments such as that shown in panel D (*n*=16-17 neurons from ≥4 cultures prepared from two rats). **(G)** Immunofluorescence microscopy showing LC3 staining in 40-μm distal axonal segments from control and DMXL2-KD DIV7 hippocampal neurons. **(H)** Quantification of total number of LC3-positive vesicles in control and DMXL2-KD neurons from experiments such as that shown in panel G (n=11 neurons from ≥3 cultures prepared from two rats). **(I)** Single frame of live microscopy showing Magic Red staining in the somatodendritic domain of control and DMXL2-KD DIV7 rat hippocampal neurons. **(J)** Quantification of Magic Red fluorescence mean intensity within the soma of control DMXL2-KD neurons from experiments such as that shown in panel I (n=17 neurons from ≥4 cultures prepared from two rats). **(K)** Immunofluorescence microscopy showing LC3 staining in the somatodendritic domain of control and DMXL2-KD DIV7 rat hippocampal neurons. **(L)** Quantification of mean LC3 intensity within the soma of control DMXL2-KD neurons from experiments such as that shown in panel K (n=18 neurons from ≥4 cultures prepared from two rats). The axonal segments in panel A and D represent mid-axon located approximately 40-200μm from the soma. All quantitative data are represented as the mean ± SD. Statistical significance was calculated using the Mann-Whitney test for panels B, E, H, J and L, and two-way ANOVA with Sidak’s multiple comparisons test for panels C and F. Actual *P* values are indicated in the figure. Scale bars: 5 μm.

## Discussion

Lysosomes are not uniformly distributed or functionally homogenous within the cells. Rather, their positioning, luminal acidity, and degradative potential are dynamically tuned by the local metabolic environment and organelle interactions (Ballabio & Bonifacino, 2020; Barral et al., 2022; Bussi & Gutierrez, 2024). Although v-ATPase-mediated proton translocation is essential for maintaining lysosomal acidification and homeostasis, the spatial coordination of v-ATPase assembly in relation to lysosome positioning and whether this regulation contributes to gradients of lysosomal vesicle pH along axons have remained largely unexplored. The present study reveals that the two structural domains of the v-ATPase, V_0_ and V_1_, display distinct subcellular distributions in non-neuronal cells and differential transport dynamics within axons, indicating a spatially regulated association of these subcomplexes. Moreover, we identify DMXL2, a core component of the mRAVE complex, as a key regulator that links V_1_–V_0_ assembly to the acidification, transport, and functional maturation of lysosomal vesicles in axons.

Because only the assembled V_1_–V_0_ complex is active, our findings provide a mechanistic basis for the heterogeneity in lysosomal vesicle acidification observed in both non-polarized cells and neurons. In non-neuronal cells, the higher concentration of V_1_ associated with V_0_ within perinuclear region likely accounts for the previously reported greater acidification of perinuclear compared to peripheral lysosomes (Johnson et al., 2016; Leung et al., 2019). Neurons present an even more striking model of spatial organization, where lysosomal vesicles must travel long distances to maintain degradative and signaling capacity across axons and dendrites (Ferguson, 2018, 2019; Roney et al., 2022). Our analyses in hippocampal neurons revealed that axonal lysosomal vesicles can be divided into at least two populations based on v-ATPase composition: vesicles containing only the V_0_ domain and vesicles containing both V_0_ and V_1_ domains. Furthermore, we found that V_0_-positive vesicles exhibit bidirectional transport, whereas V_1_-positive vesicles move almost exclusively in the retrograde direction. Moreover, co-imaging experiments revealed that V_1_–V_0_ co-localization was markedly higher in retrograde-moving vesicles than in anterograde ones. This spatially regulated assembly suggests that the distal axon functions as a site for V_1_ recruitment and holoenzyme activation, coinciding with the transition of biosynthetic or precursor vesicles into more mature lysosomal vesicles. Such compartmentalized control of v-ATPase assembly provides a mechanistic explanation for the progressive acidification of lysosomal vesicles observed along axons from distal to proximal regions, as well as for the greater acidification of retrograde compared to anterograde lysosomal vesicles (Gowrishankar et al., 2021; Gowrishankar et al., 2017; Lie et al., 2021; Overly et al., 1995).

Using SiR-Lysosome and LysoTracker as functional probes, we found that both anterograde- and retrograde-moving vesicles in the axon contained cathepsin D, consistent with their at least partial lysosomal identity. However, only the retrograde subpopulation exhibited LysoTracker fluorescence, indicative of the acidic pH required for lysosomal hydrolase activity. Indeed, only the retrograde subpopulation was stained with Magic Red cathepsin B, reflecting hydrolase activation upon acidification. These results demonstrated that the presence of lysosomal hydrolases in addition to lysosomal membrane proteins like LAMP1 in anterograde vesicles is not synonymous with degradative activity; rather, this activity requires V_1_–V_0_ assembly and v-ATPase activation in the distal axon. The observation that V_0_-only vesicles are non-acidic, yet cathepsin D-positive supports the concept of lysosomal vesicle maturation as a stepwise process beginning with delivery of hydrolases and transmembrane components via anterograde transport, followed by local V_1_–V_0_ assembly and acidification during retrograde transport along axons. In addition to maturation from anterograde lysosomal vesicles, the acquisition of endocytic or autophagic cargo may also contribute to the emergence of functionally distinct retrograde lysosomal vesicles, consistent with prior proposals that these compartments correspond to late endosomes, autolysosomes, and/or amphisomes.

Our identification of DMXL2 as a key regulator of V_1_–V_0_ association adds mechanistic significance to the emerging role of the mRAVE complex in neuronal physiology. Originally described in yeast, the RAVE complex promotes V_1_ association with V_0_ upon nutrient repletion (Smardon et al., 2002). Its mammalian homolog, mRAVE, composed of DMXL1/2, WDR7, and ROGDI, coordinates v-ATPase assembly with endolysosomal signaling and synaptic function (Jaskolka et al., 2021; Nardone et al., 2025; Peng et al., 2024). Furthermore, recent work has shown that DMXL1/2 promotes recruitment of V_1_ to lysosomes following TRPML1 activation (Lee et al., 2025). Here, we have shown that depletion of DMXL2, the major neuronal DMXL isoform, reduces the membrane association of V_1_, decreases the population of retrograde V_1_–V_0_ vesicles, and markedly impairs lysosomal vesicle acidification and cathepsin B activity in the axon. Notably, DMXL2 is enriched at synapses (Dittrich et al., 2023; Nagano et al., 2002), providing a plausible mechanism for local V_1_–V_0_ assembly in the distal axon. Loss of DMXL2 leads to an accumulation of autophagic vesicles in distal axons, as shown by the increased staining for the autophagic marker LC3B. This could be due to an inability of the less acidic lysosomes to fuse with autophagosomes, to degrade LC3B and other autophagic cargos, and/or to promote autolysosome/amphisome transport toward the soma (Lee et al., 2011; Maday et al., 2012). Thus, by orchestrating V_1_–V_0_ association, DMXL2, and by extension the mRAVE complex, ensures the spatial precision and temporal control of lysosomal vesicle activation required for axonal homeostasis.

The observation that V_1_–V_0_ assembly promotes retrograde lysosomal movement raises intriguing questions about the coupling between organelle identity and transport direction. Mature, acidified lysosomal vesicles, as well as autolysosomes/amphisomes, are known to associate with dynein-dynactin motors through adaptors such as JIP3 and JIP4 for movement toward the soma (Cason & Holzbaur, 2023; Cheng et al., 2015; Gowrishankar et al., 2021; Gowrishankar et al., 2017; Kumar et al., 2022), whereas biosynthetic/immature lysosomal vesicles preferentially engage kinesins through BORC and ARL8 for anterograde transport (De Pace et al., 2020; Farias et al., 2017; Vukoja et al., 2018). The acquisition of V_1_ and consequent acidification might thus act as a molecular cue that switches motor association, biasing vesicles toward retrograde dynein-dynactin-driven transport. Furthermore, the Rab-interacting lysosomal protein (RILP), which serves as a bridge between RAB7 and dynein, is known to stabilize v-ATPase assembly by interacting with the V_1_G1 subunit (De Luca et al., 2014), providing an additional mechanistic link between retrograde transport and maturation of lysosomal vesicles in the axon.

Our findings may provide insights into the pathogenesis of neurological disorders linked to defects in the v-ATPase and its assembly factors (Colacurcio & Nixon, 2016; Nixon & Rubinsztein, 2024). Mutations in v-ATPase subunits, including the V_0_a1 (Aoto et al., 2021; Bott et al., 2021) and V_1_B2 subunits (Yuan et al., 2014) examined in this study, are associated with a wide spectrum of neurodevelopmental and neurodegenerative disorders characterized by epilepsy, developmental delay, intellectual disability, spasticity, and/or Parkinsonism (Colacurcio & Nixon, 2016; Nixon & Rubinsztein, 2024). Mutations in subunits of the mRAVE complex cause similar neurodevelopmental phenotypes, exemplified by DMXL2 mutations in Ohtahara syndrome (Esposito et al., 2019) and ROGDI mutations in Kohlschutter-Tonz Syndrome (Mory et al., 2012; Schossig et al., 2012). Moreover, lysosomal acidification defects contribute to the pathogenesis of common neurodegenerative disorders, including Alzheimer’s disease, Parkinson’s disease, and frontotemporal dementia (Nixon & Rubinsztein, 2024). It is thus plausible that the axonal acidification, transport, and autophagic defects described in our study underlie the pathogenesis of these diseases.

In conclusion, our study uncovers a spatially orchestrated mechanism of lysosomal vesicle acidification in neuronal axons governed by v-ATPase assembly dynamics. In this model, V_0_-containing vesicles move anterogradely to deliver lysosomal transmembrane and luminal proteins to distal axonal regions, whereas V_1_–V_0_ -containing vesicles, representing acidified lysosomal vesicles, contribute to autophagosome maturation while moving retrogradely towards the soma (Fig. 7). The assembly of these domains into a functional holoenzyme is driven by the mRAVE complex in the distal axon. Loss of mRAVE disrupts V_1_ recruitment, impairs retrograde lysosomal transport, and diminishes lysosomal activity. Understanding how this assembly-transport axis is modulated under physiological and pathological conditions may open new avenues for targeting lysosomal dysfunction in neurodevelopmental and neurodegenerative diseases.

**Figure 7.**
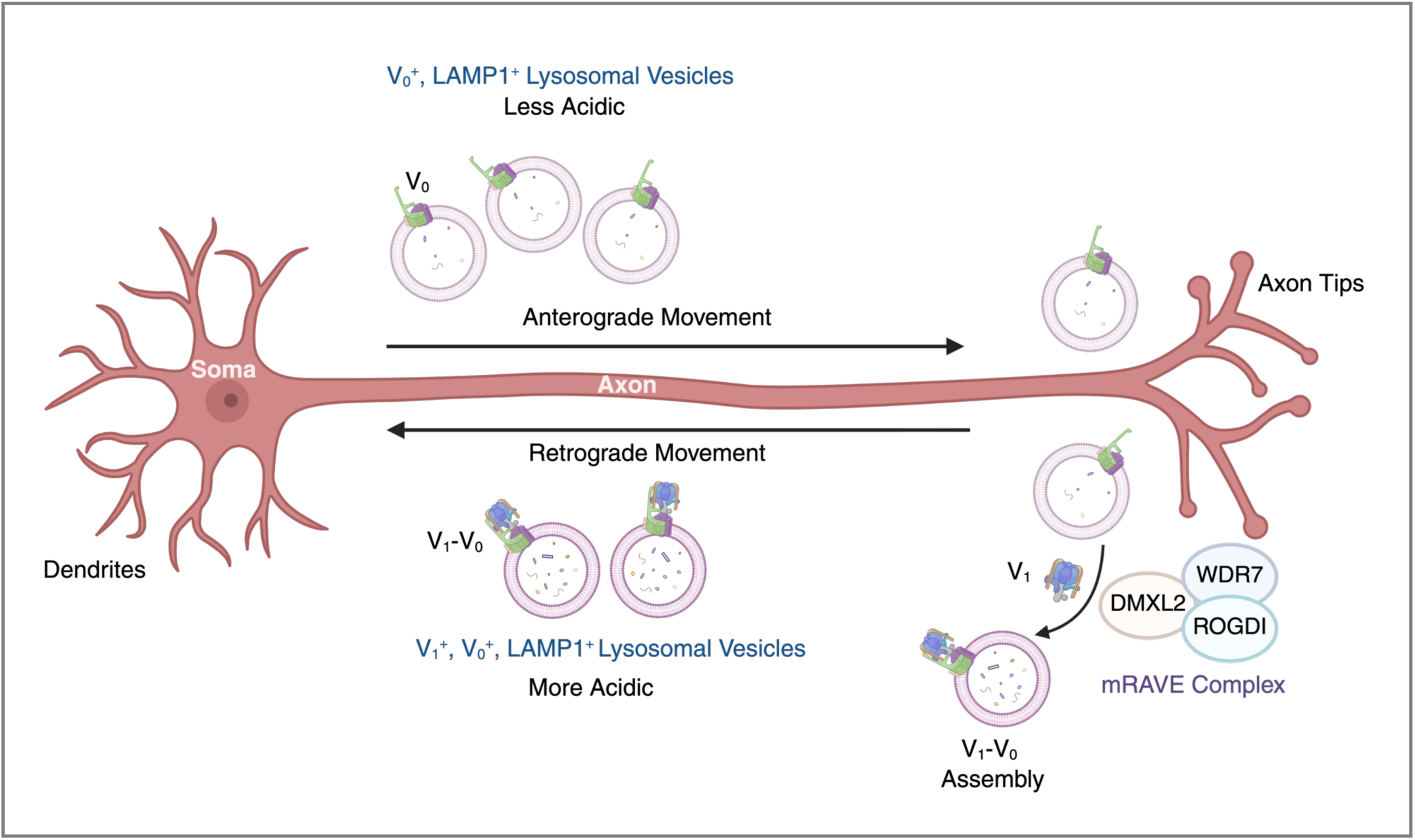
Schematic representation of how mRAVE-dependent V_1_–V_0_ assembly regulates the pH of axonal lysosomal vesicles. We propose that axonal anterograde lysosomal vesicles are less acidic because they have V_0_ but not V_1_. Upon reaching the distal axon, the mRAVE complex, composed of DMXL2, WDR7 and ROGDI, promotes recruitment of V_1_ to V_0_ domain to form an active v-ATPase holoenzyme capable of acidifying the lysosomal vesicles. This enables the retrograde transport of acidic, V_1_–V_0_-containing vesicles toward the soma.

## Materials and methods

### Antibodies

Primary antibodies: mouse anti-DMXL2 (Proteintech, Cat. 66891-2-Ig, 1:1,000 for IB), rabbit anti-ATP6V0A1 (Novus Biologicals, Cat. NBP1-89342, 1:1,000 for IB), rabbit anti-ATP6V1B2 (Abcam, Cat. ab73404, 1:1,000 for IB), mouse anti-human LAMP1 (DSHB, Cat. H4A3, 1:1,000 for IF), mouse anti-rat LAMP1 (6H2, generated in-house, 1:1 culture supernatant for IF), chicken anti-GFP (Invitrogen, Cat. A10262, 1:1,000 for IF), rat anti-mCherry (Invitrogen, Cat. M11217, 1:500 for IF), goat anti-ankyrin G (Santa Cruz Biotechnology, Cat. sc-31778, 1:100 for IF), rabbit anti-LC3 (Cell Signaling, Cat. 3868, 1:200 for IF) and anti-pan-neurofascin (extracellular) antibody (A12/18) (Antibodies Inc, Cat. 74-172) labeled with the mix-n-Stain Neurofascin CF647 Antibody Labeling Kit (Biotium, Cat. 92238), according to the manufacturer’s protocol (to label the axon initial segment for live-cell imaging). Secondary antibodies: HRP-conjugated mouse anti-α-tubulin (Santa Cruz Biotechnology, Cat. Sc-32293, 1:2,000 for IB), HRP-conjugated donkey anti-mouse IgG (Jackson ImmunoResearch, Cat. 715-035-150, 1:5,000 for IB), HRP-conjugated goat anti-rabbit IgG (Jackson ImmunoResearch, Cat. 111-035-144, 1:5,000 for IB), Alexa fluor 405 donkey anti-goat IgG (Abcam, Cat. 175664, 1:1,000 for IF), Alexa fluor 488 donkey anti-chicken IgY (Invitrogen, Cat. A78948, 1:1,000 for IF), Alexa fluor 555 donkey anti-mouse IgG (Invitrogen, Cat. A-31570, 1:1,000 for IF), Alexa fluor 555 donkey anti-rabbit IgG (Invitrogen, Cat. A-31572, 1:1,000 for IF), and Alexa fluor 568 donkey anti-rat IgG (Invitrogen, Cat. A78946, 1:1,000 for IF).

### Plasmids

The v-ATPase constructs V_0_a1–eGFP (in pGFP-N1 vector, CMV promoter) and AcGFP–V_1_B2 (in pAcGFP-C1, CMV promoter) were a kind gift from Dr. Jürgen Klingauf (Institut für Medizinische Physik und Biophysik, Universität Münster) (Bodzeta et al., 2017). The V_0_a1–mScarlet3 construct was obtained from VectorBuilder and is provided in a pRP[Exp] mammalian expression vector driven by an EF1α promoter (Vector builder, Cat. VB241010-1361tec). The mNeonGreen–V_1_B2 fusion was generated by subcloning V_1_B2 cloning into a mammalian expression vector based on the mEGFP-C1 backbone encoding the NeonGreen fluorescent protein under the CMV promoter (Addgene, Cat. 197924). The mCherry–V_1_B2 construct was generated by subcloning V_1_B2 into the pmCherry-C1 mammalian expression vector under a CMV promoter (Takara, Cat. 632524).

### Generation of stable cell lines and co-localization analysis

Stable HeLa and U2OS cell lines expressing V_0_a1–eGFP or AcGFP–V_1_B2 were generated by plasmid transfection followed by antibiotic selection and fluorescence-based enrichment. Cells were transfected at 60-70% confluence using Lipofectamine 2000 (Invitrogen, Cat. 11668027) according to the manufacturer’s instructions and allowed to recover for 18-24 h before initiating selection with 700 μg/ml geneticin (Gibco, Cat. 10131035). Resistant cells were maintained under selection for 7-10 days, then expanded and subjected to FACS to isolate GFP-positive cells. These cells were single-cell cloned in 96-well plates containing complete Dulbecco’s Modified Eagle Medium (DMEM, Quality Biologicals, Cat. 112-319-101), expanded for 2-3 weeks, and screened for robust expression and stable growth. Verified clones were then imaged to examine the localization of V_0_a1–eGFP or AcGFP–V_1_B2. For peripheral and perinuclear co-localization analysis, regions of interest (ROI) were manually defined in Fiji (ImageJ v1.54). A perinuclear ROI was drawn around the cell center encompassing the nucleus, and a peripheral ROI was drawn near the cell boundary; ROIs of equal area were selected for each cell. Pearson’s correlation coefficients were calculated using the PSC co-localization plugin in Fiji.

### Transfection of neurons

All rat procedures were conducted following the National Institutes of Health Guide for the Care and Use of Laboratory Animals, under protocol 25-011 that received ethical approval by the Eunice Kennedy Shriver National Institute of Child Health and Human Development (NICHD) Animal Care and Use Committee. Primary hippocampal neurons were isolated from embryonic day 18 (E18) rats following the protocol described earlier (Farias et al., 2017). Neurons were cultured on 18 mm glass coverslips (EMS, Cat. 72222-01) pre-coated with poly-D-lysine (Sigma, Cat. P2636), and laminin (Roche, Cat. 11243217001). Cells were placed in adhesion medium consisting of DMEM (Gibco, Cat. 2106-029), supplemented with 10 % horse serum (Gibco, Cat. 26050-088), and 100 U/mL penicillin-streptomycin (Gibco, Cat. 15070-063). After 4 h, the adhesion medium was replaced with complete Neurobasal medium consisting of Neurobasal medium (NB, Gibco, Cat. 21103049), supplemented with serum-free 1X B27 (Gibco, Cat. 17504044), 1X GlutaMax (Gibco, Cat. 35050061), and 100 U/mL penicillin-streptomycin. Neurons were transfected at DIV5 using Lipofectamine 2000. For each coverslip, 0.5-2 μg plasmid DNA was mixed with 1.1 μL Lipofectamine 2000 in 200 μL NB medium, and incubated for 20 min at room temperature to allow complex formation. Meanwhile, the culture medium was gently removed from the neurons and kept at 37 °C for re-addition after transfection (conditioned medium), and cells were overlaid with warm incomplete NB medium (Neurobasal medium without any supplement). After complex formation, the DNA-lipid mixture was added to the cells and incubated for 1 h. Neurons were then washed twice with warm NB medium. Finally, the conditioned medium was restored to each well and brought to a final volume of 1 mL with freshly prepared complete NB medium. Cells were allowed to express the constructs for 48 h prior to imaging or fixation.

### shRNA transfection

Lentiviral particles expressing *Dmxl2* shRNA were generated in HEK293T cells by co-transfecting two *Dmxl2* shRNA transfer plasmids (Vector Builder, Cat. VB250715-1369xfp and Cat. VB250715-1373xvs) with the packaging plasmids psPAX2 (Addgene, Cat. 12260), pMD2.G (Addgene, Cat. 12259), and pAdVantage (Promega, Cat. E1711). Viral supernatants were collected, filtered, and applied to cultured hippocampal neurons at DIV4. After 16-18 h, the viral medium was replaced with conditioned medium, and cultures were maintained until DIV7 for downstream processing. To visualize V_0_ and V_1_ vesicle transport live in neurons, V_0_a1–mScarlet3 or mCherry–V_1_B2 were co-transfected with either the control U6-shRNA vector or *Dmxl2*-U6-shRNA plasmid at DIV4 and imaged at DIV7. For *Dmxl2*, two different sequences were targeted: 5’-GGTCTCCTGATGGTGAATATT-3’ and 5’-CCTGGTTGGCACTGCATTTAA-3’.

### Live-cell imaging

Rat hippocampal neurons in primary culture were transfected at DIV5 and imaged at DIV7 on a Nikon Eclipse Ti spinning-disk microscope equipped with a humidified environmental chamber maintained at 37 °C and 5% CO_2_. Images were captured using a Plan Apo VC 60x/1.4 NA oil-immersion objective and an EM charged-coupled device camera (iXon DU897 from Andor). Image acquisition was controlled using NIS-Element AR software (Nikon). To identify the axon, neurons were briefly stained (1 min) with CF640R-conjugated anti-neurofascin, an axon initial segment (AIS) marker (Farias et al., 2015). V_0_a1–eGFP or V_0_a1–mScarlet3 expression plasmids were used to track V_0_ vesicles, and mCherry–V_1_B2, AcGFP–V_1_B2, or mNeonGreen–V_1_B2 expression plasmids for V_1_ vesicles. For single-color live imaging, neurons were excited with the laser channel corresponding to the fluorescently tagged protein of interest for 100–200 ms, and images were acquired every 1 s for a total duration of 180 s. For dual-color imaging, neurons in each fluorescent channel were sequentially excited for 100–200 ms and imaged every 1 s for 60–180 s.

### Live-cell lysosomal activity and labeling assays

SiR-Lysosome (Cytoskeleton, Cat. CY-SC012) was used to visualize lysosomal compartments enriched in Cathepsin D. 1 μl of the stock solution was added to 1 mL of culture medium, and cells were incubated for 20 min at 37 °C. LysoTracker Green DND-26 (Invitrogen, Cat. L7526) and LysoTracker Deep Red (Invitrogen, Cat. L12492) were used to stain acidic lysosomal compartments. Both dyes were supplied as 1 mM stocks in DMSO and diluted 1:10 to generate a 100 μM intermediate solution. From these, 1 μl was added to 1 mL of medium to obtain a final concentration of 100 nM. Cells were incubated for 10 min at 37°C. Magic Red cathepsin B substrate (Immunochemistry Technologies, Cat. SKU 937) was used to detect active Cathepsin B. The substrate was reconstituted in 50 μL DMSO and diluted 1:10 in distilled water to prepare the working solution, and 10 μl of this solution was added to 1 mL of culture medium. Cells were incubated with Magic Red for 20 min at 37 °C. After each labeling step, cells were washed with warm PBS, returned to conditioned medium, and immediately imaged using appropriate excitation wavelengths (SiR-Lysosome: 647 nm; LysoTracker Green: 488 nm; LysoTracker Deep Red: 647 nm; Magic Red Cathepsin B substrate: 561 nm).

### Kymograph analysis

Kymographs analysis was performed on time-lapse image stacks by first selecting a rectangular region of interest that encompassed a 30-μm axonal segment. Then, time-lapse image stacks were re-sliced to generate space-time plots, followed by group maximum-intensity Z-projection to produce kymographs. Vesicle trajectories were manually quantified: anterograde and retrograde movements appeared as diagonal traces with negative and positive slopes, respectively, whereas stationary vesicles appeared as vertical lines. Analysis was performed on raw images without filtering.

### Immunofluorescence microscopy

Cells were fixed in 4% paraformaldehyde (Thermo Scientific, Cat. 28906) in PBS for 15 min at room temperature. Following fixation, cells were washed twice with 1X PBS and permeabilized/blocked with immunofluorescence (IF) buffer (1X PBS, 1% BSA, and 0.1% saponin). Cells were then incubated with primary antibodies diluted in IF buffer for 1 h at 37 °C, washed twice with IF buffer, and subsequently incubated with secondary antibodies diluted in the same buffer. After secondary antibody staining, cells were washed twice with IF buffer and finally with 1X PBS. Coverslips were mounted using Fluoromount-G (EMS, Cat. 17984-25) or DAPI-Fluoromount-G (EMS, Cat. 17984-24). Confocal images were acquired on a Zeiss LSM 880 confocal microscope equipped with a Plan-Apochromat 63x/1.40 NA oil-immersion objective using the manufacturer’s recommended settings.

### Neuronal membrane extracts

DIV7 rat hippocampal neurons were washed twice with ice-cold PBS to remove serum. Cells were then incubated with homogenization buffer (0.25 M sucrose, 1 mM EDTA, and 20 mM HEPES, pH 7.4) supplemented with protease inhibitor cocktail and gently scraped into fresh microcentrifuge tubes. The suspension was homogenized by seven passes through a 20-gauge needle, followed by five passes through a 25-gauge needle. The homogenate was centrifuged at 500 x *g* for 10 min at 4 °C, and the resulting supernatant was transferred to a new tube. Iodixanol-sucrose buffered solution was prepared by mixing the homogenization medium and 50% of the iodixanol-containing working solution (OptiPrep, Stem Cell Technologies, Cat. 07820). The supernatant was loaded at the bottom of the ultracentrifuge tubes and overlaid sequentially with 1-ml layers of 50%, 40%, 35%, 30%, 28%, 24%, 20%, and 10% iodixanol-sucrose solutions. The tubes were ultracentrifuged at 120,000 x g for 18 h using a SWA1–Ti rotor. Membrane fractions were collected from the upper three layers (10%, 20% and 24%), pooled and concentrated using an ultra 30-kDa centrifugal filter unit (Millipore, Cat. UFC903008).

### Immunoblotting

Neurons were washed with ice-cold 1X PBS and lysed in 1X Laemmli buffer (Bio-Rad, Cat. 1610747) supplemented with 5% 2-mercaptoethanol (Millipore Sigma, Cat. M3148) for 15 min at room temperature. Lysates were heated at 95 °C for 5 min with gentle agitation and separated by SDS-PAGE using 4-20% precast polyacrylamide gels (Bio-Rad, Cat. 4561094). Proteins were transferred onto nitrocellulose membranes, which were then blocked for 1 h in blocking buffer containing 5% non-fat milk prepared in Tris-buffered saline with 0.01% Tween-20 (TBST; Millipore Sigma, Cat. P1379). Membranes were incubated overnight at 4 °C with primary antibodies diluted in blocking buffer with gentle rocking. Blots were washed three times (5 min each) in 1X TBST and incubated for 1 h with the appropriate secondary antibodies diluted in blocking buffer. After three additional washes in 1X TBST, bands were developed using SuperSignal West Dura extended duration substrate (Thermo Scientific, Cat. 34076) and imaged using a ChemiDoc MP imaging system (Bio-Rad, Cat. 12003154).

### Statistical analysis

All bar graphs represent the mean ± SD from multiple determinations. The total number of cells analyzed is specified in the corresponding figure legends. Statistical differences between two groups were assessed using Welch’s t-test or the Mann-Whitney test, as appropriate. For cell datasets involving repeated measures across multiple conditions, paired Friedman tests with Dunn’s post hoc correction were used. Comparisons among multiple groups were performed using two-way ANOVA followed by Tukey’s or Sidak’s multiple comparison tests. All analyses were conducted using Graph Pad Prism 9.

### Data availability

Data are available in the article itself and its supplementary materials, or otherwise available upon request.

## Acknowledgements

We thank Jürgen Klingauf for the kind gift of reagents. This work was supported by the Intramural Program of the NICHD (ZIA HD001607). We thank Dr. Ganesh Shelke for guidance on neuronal membrane extraction.

## Author contributions

Conceptualization, S. Verma and J. S. Bonifacino; Methodology, S. Verma, N. A. Tirumala, X. Zhu, R. De Pace; Investigation, S. Verma; Writing, S. Verma and J. S. Bonifacino; Funding acquisition, J. S. Bonifacino; Supervision, J. S. Bonifacino.

## Abbreviation

AcGFP: *Aequorea coerulescens* green fluorescent protein
AIS: Axon initial segment
AnkG: Ankyrin G
ATP: Adenosine triphosphate
BSA: Bovine serum albumin
BORC: BLOC-one-related complex
CatB: Cathepsin B
CMV: Cytomegalovirus (promoter)
DIV: Days in vitro
DMEM: Dulbecco’s Modified Eagle Medium
DMXL1/2: Dmx-like protein 1 / 2
DMSO: Dimethyl sulfoxide
eGFP: Enhanced green fluorescent protein
FACS: Fluorescence-activated cell sorting
GFP: Green fluorescent protein
HRP: Horseradish peroxidase
IB: Immunoblot
IF: Immunofluorescence
KD: Knockdown
LAMP1: Lysosome-associated membrane protein 1
LC3: Microtubule-associated protein 1 light chain 3
mRAVE: Metazoan regulator of the ATPase of vacuolar and endosomal membranes
NB: Neurobasal medium
PBS: Phosphate-buffered saline
ROI: Region of interest
RAVE: Regulator of the ATPase of vacuolar and endosomal membranes
RILP: Rab-interacting lysosomal protein
shRNA: Short hairpin RNA
SiR: Silicon–rhodamine
TBST: Tris-buffered saline with Tween-20
TRPML1: Transient receptor potential mucolipin 1
U2OS: Human osteosarcoma cell line
V-ATPase: Vacuolar-type H⁺-ATPase
V_0_: Membrane-embedded domain of the v-ATPase
V_1_: Peripheral, ATP-hydrolyzing domain of the v-ATPase
WT: Wild type

---

**Figure S1.**
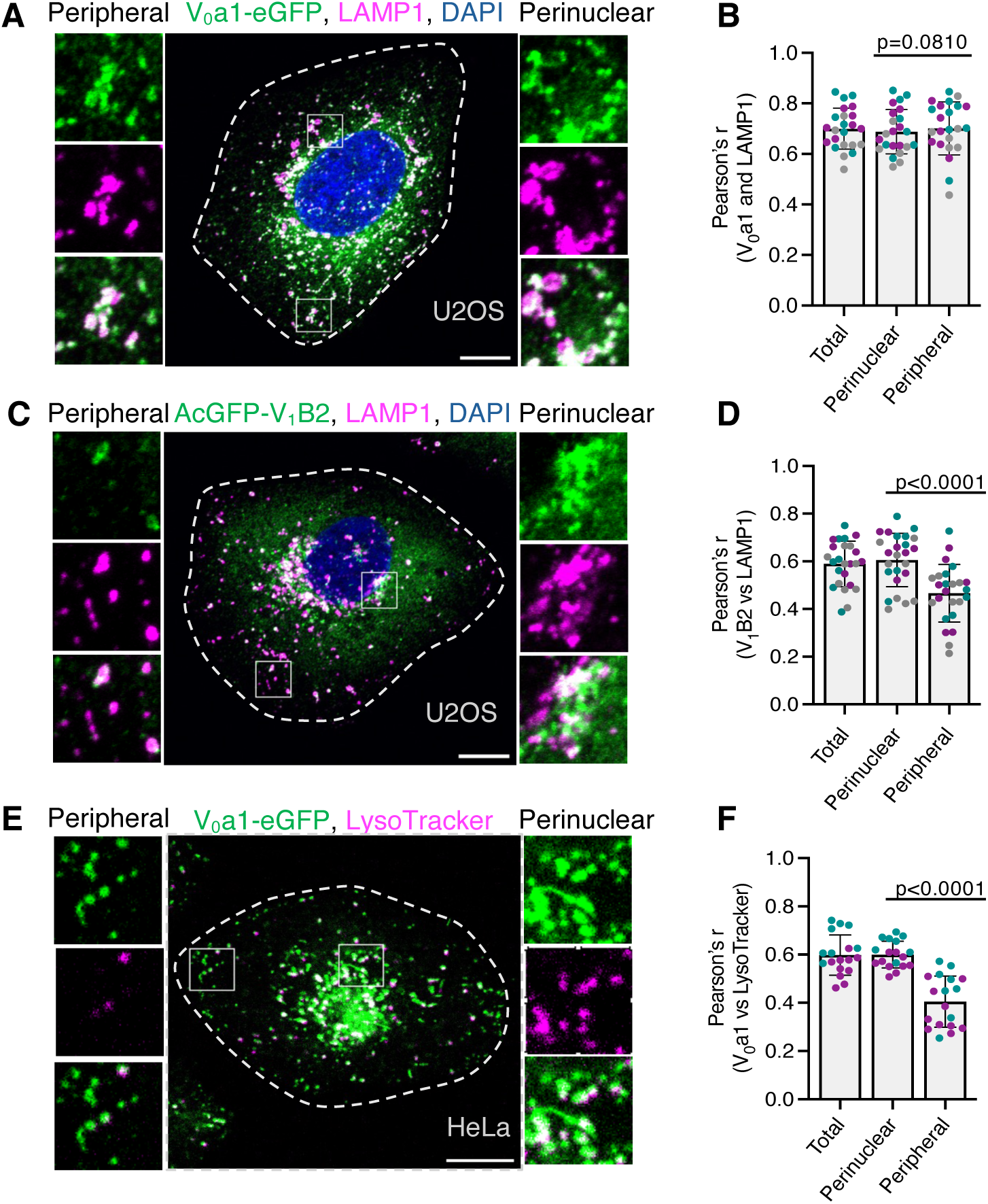
Co-localization of v-ATPase subunits V_0_a1 and V_1_B2 with lysosomal markers in HeLa and U2OS cells. **(A,C)** Immunofluorescence microscopy showing co-localization of stably expressed V_0_a1–eGFP (green) (A) or AcGFP–V_1_B2 (green) (C) with endogenous LAMP1 (magenta) in U2OS cells. Nuclei were stained with DAPI (blue). Magnified views of the boxed areas in the peripheral and perinuclear regions are shown at left and right, respectively. **(B,D)** Quantification of the co-localization between LAMP1 and V_0_a1–eGFP (B) or AcGFP–V_1_B2 (D) in total, perinuclear, and peripheral regions, expressed as Pearson’s correlation coefficients (Pearson’s r), from experiments such as those shown in panels A and C (*n*=23-25 cells from three independent experiments). **(E)** Single frame live-cell images of HeLa cells showing the co-localization of stably expressed V_0_a1–eGFP (green) with LysoTracker (magenta). Magnified views of the boxed areas in the peripheral and perinuclear regions are shown at left and right, respectively**. (F)** Quantification of the co-localization between LysoTracker and V_0_a1–eGFP in total, perinuclear, and peripheral regions, expressed as Pearson’s r, from experiments such as those shown in panel E (*n*=18-19 cells from two independent experiments). All quantitative data are represented as the mean ± SD. Statistical significance was assessed using the Friedman test with Dunn’s multiple comparisons test. Actual *P* values are indicated in the figure. Scale bars: 10 μm.

**Figure S2.**
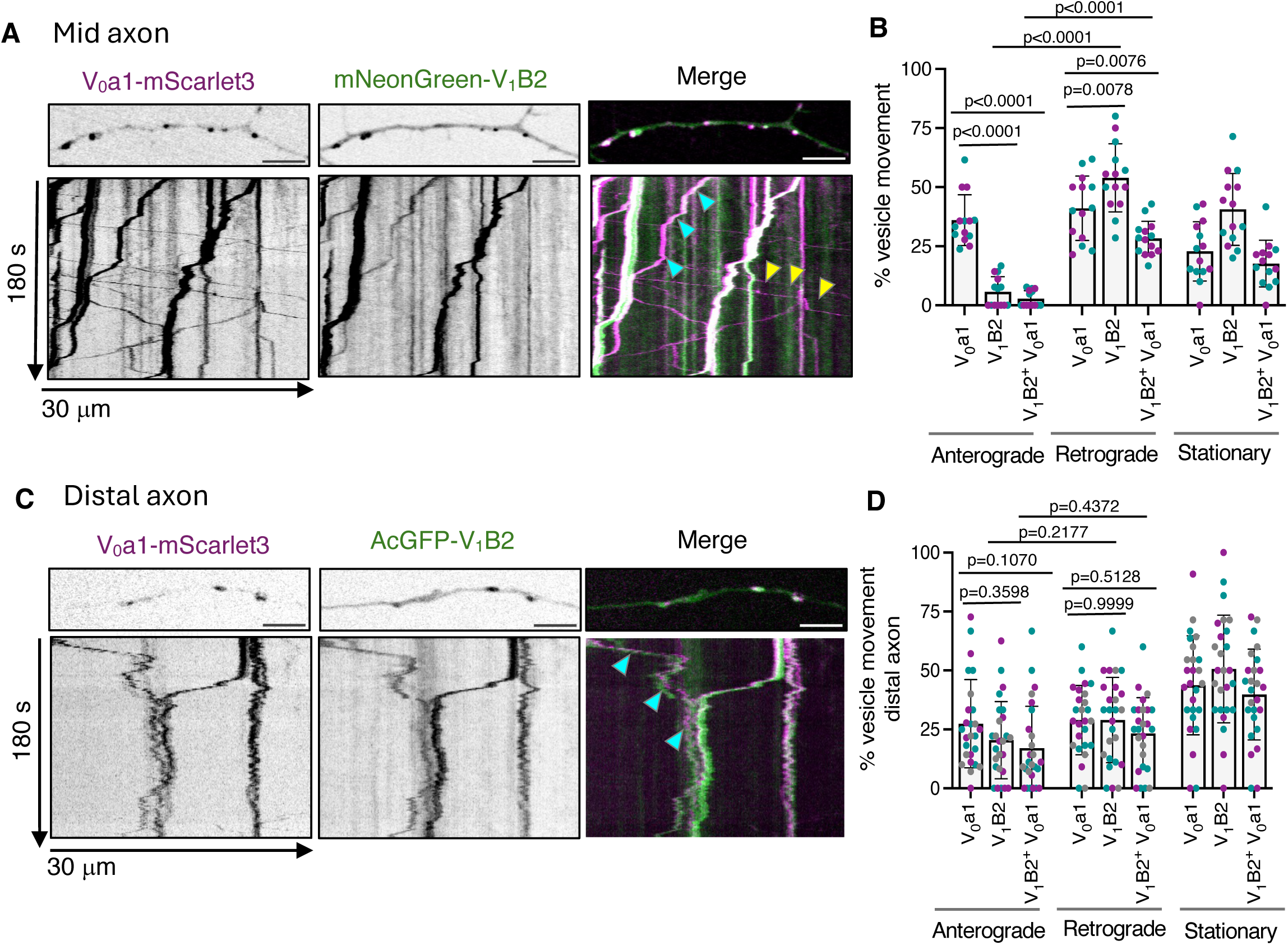
Transport dynamics of vesicles containing V_0_a1 and V_1_B2 subunits in rat hippocampal neurons. **(A)** Single frames (top) and corresponding kymographs (bottom) of 30-μm axonal segments located approximately 50 μm from the soma of DIV7 rat hippocampal neurons co-expressing V_0_a1–mScarlet3 and mNeonGreen–V_1_B2 imaged live for 180 s. **(B)** Quantification of the proportion of anterograde, retrograde, and stationary vesicles in axons from neurons co-expressing V_0_a1–mScarlet3 and mNeonGreen–V_1_B2 from experiments such as that shown in panel A (*n*=14 neurons from ζ4 cultures prepared from two rats). Cyan arrowheads indicate V_1_-V_0_-positive vesicles and yellow arrows indicate V_0_-only-positive vesicles. (**C)** Single frames (top) and corresponding kymographs (bottom) of 30-μm distal axonal segments, located approximately 20 μm from the axon tip of DIV7 rat hippocampal neurons co-expressing V_0_a1–mScarlet3 and AcGFP–V_1_B2, imaged live for 180 s. Cyan arrowheads indicate V_1_-V_0_-positive vesicles. **(D)** Quantification of the proportion of anterograde, retrograde, and stationary vesicles in axons from neurons co-expressing V_0_a1–mScarlet3 and AcGFP–V_1_B2 from experiments such as that shown in panel C (*n*=26 neurons from ζ4 cultures prepared from three rats). Statistical significance was calculated using two-way ANOVA with Tukey’s multiple comparisons test for panel F. Actual *P* values are indicated in the figure. Scale bars: 5 μm.

## Movies

**Video S1.**
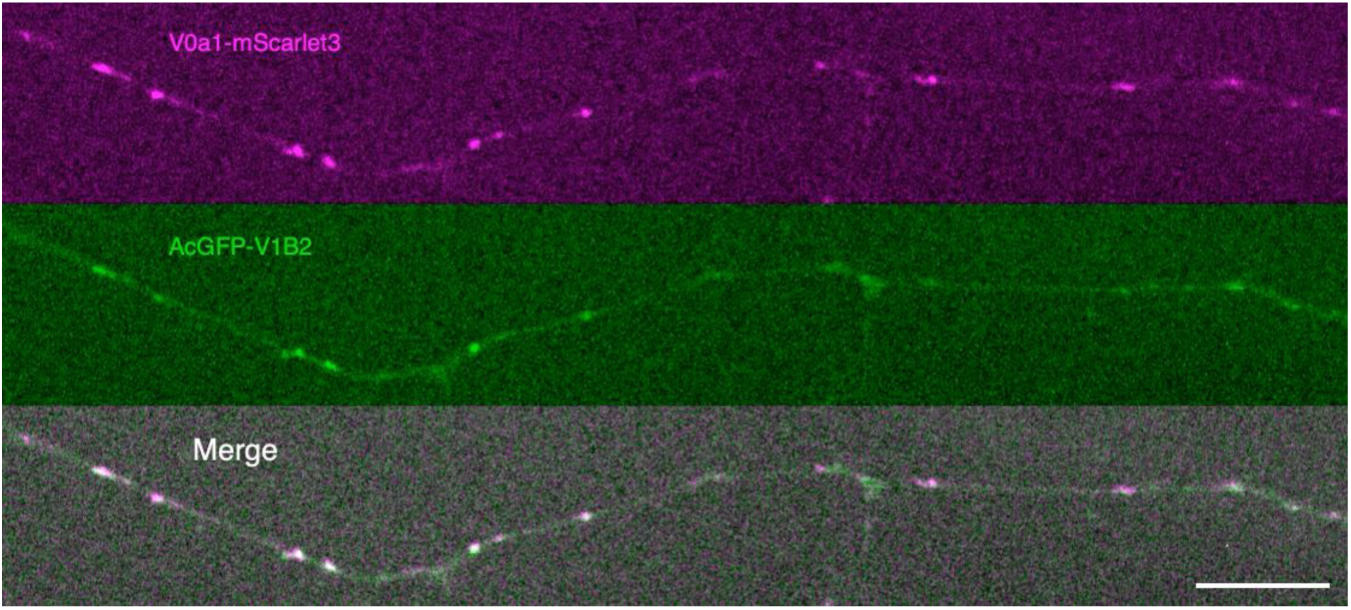
Video showing a 100-μm long section from the axon shaft of a DIV7 rat hippocampal neuron co-expressing V_0_-mScarlet3 (magenta) and AcGFP-V1B2 (green) and imaged by live-cell spinning-disk fluorescence microscopy. The time-series was recorded for 180 s with 1 s between frames. Notice the bidirectional movement of V_0_a1, retrograde movement of V_1_B2, and merged vesicles (white). Scale bar: 10 μm

**Video S2.**
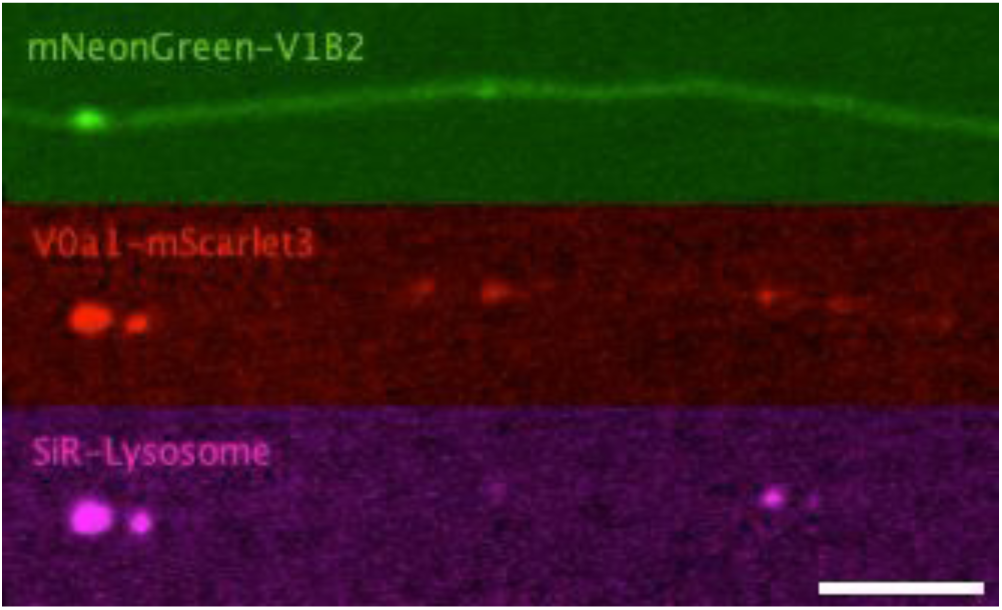
Video showing a 30-μm long section from the axon shaft of a DIV7 rat hippocampal neuron co-expressing V_0_a1-mScarlet3 (red) and mNeon-GreenV_1_B2 (green), stained with SiR-lysosome (magenta) and imaged by live-cell spinning-disk fluorescence microscopy. The time-series was recorded for 60 s at 1 s intervals. Notice the co-movement of SiR-lysosome with V_0_a1 and V_1_B2 vesicles. Scale bar: 5 μm

## References

1. Albrecht, L. V., Tejeda-Munoz, N., & De Robertis, E. M. (2020). Protocol for Probing Regulated Lysosomal Activity and Function in Living Cells. STAR Protoc, 1(3), 100132. 10.1016/j.xpro.2020.100132

2. Aoto, K., Kato, M., Akita, T., Nakashima, M., Mutoh, H., Akasaka, N., Tohyama, J., Nomura, Y., Hoshino, K., Ago, Y., Tanaka, R., Epstein, O., Ben-Haim, R., Heyman, E., Miyazaki, T., Belal, H., Takabayashi, S., Ohba, C., Takata, A.,…Saitsu, H. (2021). ATP6V0A1 encoding the a1-subunit of the V0 domain of vacuolar H(+)-ATPases is essential for brain development in humans and mice. Nat Commun, 12(1), 2107. 10.1038/s41467-021-22389-5

3. Ballabio, A., & Bonifacino, J. S. (2020). Lysosomes as dynamic regulators of cell and organismal homeostasis. Nat Rev Mol Cell Biol, 21(2), 101–118. 10.1038/s41580-019-0185-4

4. Barral, D. C., Staiano, L., Guimas Almeida, C., Cutler, D. F., Eden, E. R., Futter, C. E., Galione, A., Marques, A. R. A., Medina, D. L., Napolitano, G., Settembre, C., Vieira, O. V., Aerts, J., Atakpa-Adaji, P., Bruno, G., Capuozzo, A., De Leonibus, E., Di Malta, C., Escrevente, C.,…Seabra, M. C. (2022). Current methods to analyze lysosome morphology, positioning, motility and function. Traffic, 23(5), 238–269. 10.1111/tra.12839

5. Bodzeta, A., Kahms, M., & Klingauf, J. (2017). The Presynaptic v-ATPase Reversibly Disassembles and Thereby Modulates Exocytosis but Is Not Part of the Fusion Machinery. Cell Rep, 20(6), 1348–1359. 10.1016/j.celrep.2017.07.040

6. Bott, L. C., Forouhan, M., Lieto, M., Sala, A. J., Ellerington, R., Johnson, J. O., Speciale, A. A., Criscuolo, C., Filla, A., Chitayat, D., Alkhunaizi, E., Shannon, P., Nemeth, A. H., Italian Undiagnosed Diseases, N., Angelucci, F., Lim, W. F., Striano, P., Zara, F., Helbig, I.,…Rinaldi, C. (2021). Variants in ATP6V0A1 cause progressive myoclonus epilepsy and developmental and epileptic encephalopathy. Brain Commun, 3(4), fcab245. 10.1093/braincomms/fcab245

7. Bussi, C., & Gutierrez, M. G. (2024). One size does not fit all: Lysosomes exist in biochemically and functionally distinct states. PLoS Biol, 22(3), e3002576. 10.1371/journal.pbio.3002576

8. Calcraft, P. J., Ruas, M., Pan, Z., Cheng, X., Arredouani, A., Hao, X., Tang, J., Rietdorf, K., Teboul, L., Chuang, K. T., Lin, P., Xiao, R., Wang, C., Zhu, Y., Lin, Y., Wyatt, C. N., Parrington, J., Ma, J., Evans, A. M.,…Zhu, M. X. (2009). NAADP mobilizes calcium from acidic organelles through two-pore channels. Nature, 459(7246), 596–600. 10.1038/nature08030

9. Cang, C., Aranda, K., Seo, Y. J., Gasnier, B., & Ren, D. (2015). TMEM175 Is an Organelle K(+) Channel Regulating Lysosomal Function. Cell, 162(5), 1101–1112. 10.1016/j.cell.2015.08.002

10. Cason, S. E., & Holzbaur, E. L. F. (2023). Axonal transport of autophagosomes is regulated by dynein activators JIP3/JIP4 and ARF/RAB GTPases. J Cell Biol, 222(12). 10.1083/jcb.202301084

11. Chazotte, B. (2011). Labeling lysosomes in live cells with LysoTracker. Cold Spring Harb Protoc, 2011(2), pdb prot5571. 10.1101/pdb.prot5571

12. Cheng, X. T., Zhou, B., Lin, M. Y., Cai, Q., & Sheng, Z. H. (2015). Axonal autophagosomes recruit dynein for retrograde transport through fusion with late endosomes. J Cell Biol, 209(3), 377–386. 10.1083/jcb.201412046

13. Colacurcio, D. J., & Nixon, R. A. (2016). Disorders of lysosomal acidification-The emerging role of v-ATPase in aging and neurodegenerative disease. Ageing Res Rev, 32, 75–88. 10.1016/j.arr.2016.05.004

14. Collins, M. P., & Forgac, M. (2020). Regulation and function of V-ATPases in physiology and disease. Biochim Biophys Acta Biomembr, 1862(12), 183341. 10.1016/j.bbamem.2020.183341

15. De Luca, M., Cogli, L., Progida, C., Nisi, V., Pascolutti, R., Sigismund, S., Di Fiore, P. P., & Bucci, C. (2014). RILP regulates vacuolar ATPase through interaction with the V1G1 subunit. J Cell Sci, 127(Pt 12), 2697–2708. 10.1242/jcs.142604

16. De Pace, R., Britt, D. J., Mercurio, J., Foster, A. M., Djavaherian, L., Hoffmann, V., Abebe, D., & Bonifacino, J. S. (2020). Synaptic Vesicle Precursors and Lysosomes Are Transported by Different Mechanisms in the Axon of Mammalian Neurons. Cell Rep, 31(11), 107775. 10.1016/j.celrep.2020.107775

17. Dittrich, A., Ramesh, G., Jung, M., & Schmitz, F. (2023). Rabconnectin-3alpha/DMXL2 Is Locally Enriched at the Synaptic Ribbon of Rod Photoreceptor Synapses. Cells, 12(12). 10.3390/cells12121665

18. Esposito, A., Falace, A., Wagner, M., Gal, M., Mei, D., Conti, V., Pisano, T., Aprile, D., Cerullo, M. S., De Fusco, A., Giovedi, S., Seibt, A., Magen, D., Polster, T., Eran, A., Stenton, S. L., Fiorillo, C., Ravid, S., Mayatepek, E.,…Guerrini, R. (2019). Biallelic DMXL2 mutations impair autophagy and cause Ohtahara syndrome with progressive course. Brain, 142(12), 3876–3891. 10.1093/brain/awz326

19. Farfel-Becker, T., Roney, J. C., Cheng, X. T., Li, S., Cuddy, S. R., & Sheng, Z. H. (2019). Neuronal Soma-Derived Degradative Lysosomes Are Continuously Delivered to Distal Axons to Maintain Local Degradation Capacity. Cell Rep, 28(1), 51–64 e54. 10.1016/j.celrep.2019.06.013

20. Farias, G. G., Guardia, C. M., Britt, D. J., Guo, X., & Bonifacino, J. S. (2015). Sorting of Dendritic and Axonal Vesicles at the Pre-axonal Exclusion Zone. Cell Rep, 13(6), 1221–1232. 10.1016/j.celrep.2015.09.074

21. Farias, G. G., Guardia, C. M., De Pace, R., Britt, D. J., & Bonifacino, J. S. (2017). BORC/kinesin-1 ensemble drives polarized transport of lysosomes into the axon. Proc Natl Acad Sci U S A, 114(14), E2955–E2964. 10.1073/pnas.1616363114

22. Ferguson, S. M. (2018). Axonal transport and maturation of lysosomes. Curr Opin Neurobiol, 51, 45–51. 10.1016/j.conb.2018.02.020

23. Ferguson, S. M. (2019). Neuronal lysosomes. Neurosci Lett, 697, 1–9. 10.1016/j.neulet.2018.04.005

24. Freeman, S. A., Grinstein, S., & Orlowski, J. (2023). Determinants, maintenance, and function of organellar pH. Physiol Rev, 103(1), 515–606. 10.1152/physrev.00009.2022

25. Gowrishankar, S., Lyons, L., Rafiq, N. M., Roczniak-Ferguson, A., De Camilli, P., & Ferguson, S. M. (2021). Overlapping roles of JIP3 and JIP4 in promoting axonal transport of lysosomes in human iPSC-derived neurons. Mol Biol Cell, 32(11), 1094–1103. 10.1091/mbc.E20-06-0382

26. Gowrishankar, S., Wu, Y., & Ferguson, S. M. (2017). Impaired JIP3-dependent axonal lysosome transport promotes amyloid plaque pathology. J Cell Biol, 216(10), 3291–3305. 10.1083/jcb.201612148

27. Jaskolka, M. C., Winkley, S. R., & Kane, P. M. (2021). RAVE and Rabconnectin-3 Complexes as Signal Dependent Regulators of Organelle Acidification. Front Cell Dev Biol, 9, 698190. 10.3389/fcell.2021.698190

28. Johnson, D. E., Ostrowski, P., Jaumouille, V., & Grinstein, S. (2016). The position of lysosomes within the cell determines their luminal pH. J Cell Biol, 212(6), 677–692. 10.1083/jcb.201507112

29. Kane, P. M. (2012). Targeting reversible disassembly as a mechanism of controlling V-ATPase activity. Curr Protein Pept Sci, 13(2), 117–123. 10.2174/138920312800493142

30. Kornak, U., Kasper, D., Bosl, M. R., Kaiser, E., Schweizer, M., Schulz, A., Friedrich, W., Delling, G., & Jentsch, T. J. (2001). Loss of the ClC-7 chloride channel leads to osteopetrosis in mice and man. Cell, 104(2), 205–215. 10.1016/s0092-8674(01)00206-9

31. Kumar, G., Chawla, P., Dhiman, N., Chadha, S., Sharma, S., Sethi, K., Sharma, M., & Tuli, A. (2022). RUFY3 links Arl8b and JIP4-Dynein complex to regulate lysosome size and positioning. Nat Commun, 13(1), 1540. 10.1038/s41467-022-29077-y

32. Lawrence, R. E., & Zoncu, R. (2019). The lysosome as a cellular centre for signalling, metabolism and quality control. Nat Cell Biol, 21(2), 133–142. 10.1038/s41556-018-0244-7

33. Lee, C., Eldridge, M. J. G., Gonzalez-Lozano, M. A., Bresnahan, T., Niday, Z., Del Camino, D., Fu, T., Paulo, J. A., Moran, M. M., Helaine, S., & Harper, J. W. (2025). DMXL1 promotes recruitment of V1-ATPase to lysosomes upon TRPML1 activation. Nat Struct Mol Biol, 32(10), 2060–2075. 10.1038/s41594-025-01581-x

34. Lee, S., Sato, Y., & Nixon, R. A. (2011). Lysosomal proteolysis inhibition selectively disrupts axonal transport of degradative organelles and causes an Alzheimer’s-like axonal dystrophy. J Neurosci, 31(21), 7817–7830. 10.1523/JNEUROSCI.6412-10.2011

35. Leung, K., Chakraborty, K., Saminathan, A., & Krishnan, Y. (2019). A DNA nanomachine chemically resolves lysosomes in live cells. Nat Nanotechnol, 14(2), 176–183. 10.1038/s41565-018-0318-5

36. Li, C. H., Kersten, N., Ozkan, N., Nguyen, D. T. M., Koppers, M., Post, H., Altelaar, M., & Farias, G. G. (2024). Spatiotemporal proteomics reveals the biosynthetic lysosomal membrane protein interactome in neurons. Nat Commun, 15(1), 10829. 10.1038/s41467-024-55052-w

37. Lie, P. P. Y., Yang, D. S., Stavrides, P., Goulbourne, C. N., Zheng, P., Mohan, P. S., Cataldo, A. M., & Nixon, R. A. (2021). Post-Golgi carriers, not lysosomes, confer lysosomal properties to pre-degradative organelles in normal and dystrophic axons. Cell Rep, 35(4), 109034. 10.1016/j.celrep.2021.109034

38. Maday, S., Wallace, K. E., & Holzbaur, E. L. (2012). Autophagosomes initiate distally and mature during transport toward the cell soma in primary neurons. J Cell Biol, 196(4), 407–417. 10.1083/jcb.201106120

39. Maxson, M. E., & Grinstein, S. (2014). The vacuolar-type H(+)-ATPase at a glance - more than a proton pump. J Cell Sci, 127(Pt 23), 4987–4993. 10.1242/jcs.158550

40. Mindell, J. A. (2012). Lysosomal acidification mechanisms. Annu Rev Physiol, 74, 69–86. 10.1146/annurev-physiol-012110-142317

41. Mory, A., Dagan, E., Illi, B., Duquesnoy, P., Mordechai, S., Shahor, I., Romani, S., Hawash-Moustafa, N., Mandel, H., Valente, E. M., Amselem, S., & Gershoni-Baruch, R. (2012). A nonsense mutation in the human homolog of Drosophila rogdi causes Kohlschutter-Tonz syndrome. Am J Hum Genet, 90(4), 708–714. 10.1016/j.ajhg.2012.03.005

42. Nagano, F., Kawabe, H., Nakanishi, H., Shinohara, M., Deguchi-Tawarada, M., Takeuchi, M., Sasaki, T., & Takai, Y. (2002). Rabconnectin-3, a novel protein that binds both GDP/GTP exchange protein and GTPase-activating protein for Rab3 small G protein family. J Biol Chem, 277(12), 9629–9632. 10.1074/jbc.C100730200

43. Nardone, C., Mintseris, J., He, D., Rutter, J. C., Ebert, B. L., Gygi, S. P., & Rapoport, T. (2025). A heterotrimeric protein complex assembles the metazoan V-ATPase upon dissipation of proton gradients. Nat Struct Mol Biol, 32(10), 2076–2087. 10.1038/s41594-025-01610-9

44. Nixon, R. A., & Rubinsztein, D. C. (2024). Mechanisms of autophagy-lysosome dysfunction in neurodegenerative diseases. Nat Rev Mol Cell Biol, 25(11), 926–946. 10.1038/s41580-024-00757-5

45. Oot, R. A., Couoh-Cardel, S., Sharma, S., Stam, N. J., & Wilkens, S. (2017). Breaking up and making up: The secret life of the vacuolar H(+) -ATPase. Protein Sci, 26(5), 896–909. 10.1002/pro.3147

46. Overly, C. C., Lee, K. D., Berthiaume, E., & Hollenbeck, P. J. (1995). Quantitative measurement of intraorganelle pH in the endosomal-lysosomal pathway in neurons by using ratiometric imaging with pyranine. Proc Natl Acad Sci U S A, 92(8), 3156–3160. 10.1073/pnas.92.8.3156

47. Peng, H., Wang, L., Gao, Y., Liu, H., Lin, G., Kong, Y., Xu, P., Liu, H., Yuan, Q., Liu, H., Song, L., Yang, T., & Wu, H. (2024). DMXL2 Is Required for Endocytosis and Recycling of Synaptic Vesicles in Auditory Hair Cells. J Neurosci, 44(38). 10.1523/JNEUROSCI.1405-23.2024

48. Pu, J., Guardia, C. M., Keren-Kaplan, T., & Bonifacino, J. S. (2016). Mechanisms and functions of lysosome positioning. J Cell Sci, 129(23), 4329–4339. 10.1242/jcs.196287

49. Roney, J. C., Cheng, X. T., & Sheng, Z. H. (2022). Neuronal endolysosomal transport and lysosomal functionality in maintaining axonostasis. J Cell Biol, 221(3). 10.1083/jcb.202111077

50. Saftig, P., & Klumperman, J. (2009). Lysosome biogenesis and lysosomal membrane proteins: trafficking meets function. Nat Rev Mol Cell Biol, 10(9), 623–635. 10.1038/nrm2745

51. Schossig, A., Wolf, N. I., Fischer, C., Fischer, M., Stocker, G., Pabinger, S., Dander, A., Steiner, B., Tonz, O., Kotzot, D., Haberlandt, E., Amberger, A., Burwinkel, B., Wimmer, K., Fauth, C., Grond-Ginsbach, C., Koch, M. J., Deichmann, A., von Kalle, C.,…Zschocke, J. (2012). Mutations in ROGDI Cause Kohlschutter-Tonz Syndrome. Am J Hum Genet, 90(4), 701–707. 10.1016/j.ajhg.2012.02.012

52. Seol, J. H., Shevchenko, A., Shevchenko, A., & Deshaies, R. J. (2001). Skp1 forms multiple protein complexes, including RAVE, a regulator of V-ATPase assembly. Nat Cell Biol, 3(4), 384–391. 10.1038/35070067

53. Smardon, A. M., Tarsio, M., & Kane, P. M. (2002). The RAVE complex is essential for stable assembly of the yeast V-ATPase. J Biol Chem, 277(16), 13831–13839. 10.1074/jbc.M200682200

54. Soyombo, A. A., Tjon-Kon-Sang, S., Rbaibi, Y., Bashllari, E., Bisceglia, J., Muallem, S., & Kiselyov, K. (2006). TRP-ML1 regulates lysosomal pH and acidic lysosomal lipid hydrolytic activity. J Biol Chem, 281(11), 7294–7301. 10.1074/jbc.M508211200

55. Toei, M., Saum, R., & Forgac, M. (2010). Regulation and isoform function of the V-ATPases. Biochemistry, 49(23), 4715–4723. 10.1021/bi100397s

56. Vasanthakumar, T., & Rubinstein, J. L. (2020). Structure and Roles of V-type ATPases. Trends Biochem Sci, 45(4), 295–307. 10.1016/j.tibs.2019.12.007

57. Vukoja, A., Rey, U., Petzoldt, A. G., Ott, C., Vollweiter, D., Quentin, C., Puchkov, D., Reynolds, E., Lehmann, M., Hohensee, S., Rosa, S., Lipowsky, R., Sigrist, S. J., & Haucke, V. (2018). Presynaptic Biogenesis Requires Axonal Transport of Lysosome-Related Vesicles. Neuron, 99(6), 1216–1232 e1217. 10.1016/j.neuron.2018.08.004

58. Xu, H., & Ren, D. (2015). Lysosomal physiology. Annu Rev Physiol, 77, 57–80. 10.1146/annurev-physiol-021014-071649

59. Yuan, Y., Zhang, J., Chang, Q., Zeng, J., Xin, F., Wang, J., Zhu, Q., Wu, J., Lu, J., Guo, W., Yan, X., Jiang, H., Zhou, B., Li, Q., Gao, X., Yuan, H., Yang, S., Han, D., Mao, Z.,…Dai, P. (2014). De novo mutation in ATP6V1B2 impairs lysosome acidification and causes dominant deafness-onychodystrophy syndrome. Cell Res, 24(11), 1370–1373. 10.1038/cr.2014.77

